# The F-box cofactor UFO redirects the LEAFY floral regulator to novel *cis*-elements

**DOI:** 10.1101/2022.06.14.495942

**Authors:** Philippe Rieu, Laura Turchi, Emmanuel Thévenon, Eleftherios Zarkadas, Max Nanao, Hicham Chahtane, Gabrielle Tichtinsky, Jérémy Lucas, Romain Blanc-Mathieu, Chloe Zubieta, Guy Schoehn, François Parcy

**Affiliations:** Univ. Grenoble Alpes, CEA, CNRS, INRAE, Laboratoire Physiologie Cellulaire et Végétale, IRIG-DBSCI-LPCV, 17 avenue des martyrs, F-38054, Grenoble, France; Translational Innovation in Medicine and Complexity, Univ. Grenoble Alpes, CNRS, Rond-Point de la Croix de Vie, F-38706, Grenoble, France; Univ. Grenoble Alpes, CNRS, CEA, IBS, F-38000 Grenoble, France; Structural Biology Group, European Synchrotron Radiation Facility, 71 Avenue des Martyrs, F-38000, Grenoble, France

**Keywords:** Transcriptional complex, flower development, F-box protein, LEAFY, UFO, cryo-electron microscopy, ampDAP-seq

## Abstract

In angiosperms, flower patterning requires the localized expression of the *APETALA3* (*AP3*) floral homeotic gene involved in petal and stamen development. *AP3* is synergistically induced by the master transcription factor (TF) LEAFY (LFY) and the F-box protein UNUSUAL FLORAL ORGANS (UFO), but the molecular mechanism underlying this synergy has remained unknown. Here we show that the connection to ubiquitination pathways suggested by the F-box domain of UFO is mostly dispensable for its function and that UFO instead acts by forming a transcriptional complex with LFY and binds to newly discovered regulatory elements. Cryo-electron microscopy explains how a LFY-UFO complex forms on these novel DNA sites due to direct interaction of UFO with LFY and DNA. Finally, we show that this complex has a deep evolutionary origin, largely predating flowering plants. This work reveals a novel mechanism of an F-box protein in directly modulating the DNA-binding specificity of a master TF.

## INTRODUCTION

Angiosperm flowers are made of four types of organs (sepals, petals, stamens and carpels) arranged in concentric whorls. The patterning of flower meristems requires the localized induction of the ABCE floral homeotic genes that determine specific floral organ identities. The organ specification process is largely controlled by the master transcription factor (TF) LEAFY (LFY) that activates the floral organ homeotic genes in specific territories (Irish, 2010; Moyroud et al., 2010). LFY activates the A class gene *APETALA1* (*AP1*) uniformly in the early flower meristem (Parcy et al., 1998; Wagner et al., 1999), while other activations are local and require the activity of cofactors. LFY, in conjunction with the TF WUSCHEL, regulates the C class gene *AGAMOUS* (*AG*; Lohmann et al., 2001). The activation of the B class gene *APETALA 3 (AP3)*, requires the combined activity of LFY and the spatially-delineated cofactor UNUSUAL FLORAL ORGANS (UFO; Lee et al., 1997; Levin and Meyerowitz, 1995; Wilkinson and Haughn, 1995). In Arabidopsis, the main function of LFY and UFO is to activate *AP3* (Krizek and Meyerowitz, 1996) but in several species their joint role goes well beyond *B* genes activation and is key to floral meristem and inflorescence development (Ikeda-Kawakatsu et al., 2012; Lippman et al., 2008; Souer et al., 2008).

At the molecular level, little is known on the nature of LFY-UFO synergy. Unlike most floral regulators, *UFO* does not encode for a TF but for an F-box protein, one of the first to be described in plants (Ingram et al., 1997; Samach et al., 1999; Simon et al., 1994). UFO is part of a SKP1-Cullin1-F-box (SCF) E3 ubiquitin ligase complex in which the F-box domain of UFO directly interacts with ARABIDOPSIS SKP1-LIKE (ASK) proteins (Samach et al., 1999; Wang et al., 2003). In addition, its predicted C-terminal Kelch-type β-propeller domain physically interacts with LFY DNA Binding Domain (DBD; Chae et al., 2008) suggesting that LFY might be the target of SCF^UFO^.

We focused on the LFY-UFO interaction to understand how a component of an E3 ligase complex like UFO modulates LFY activity. As the control of TF activity through proteolytic and non-proteolytic ubiquitination is a well-described mechanism (Geng et al., 2012), UFO was previously proposed to regulate LFY activity through such post-translational modifications (Chae et al., 2008; Risseeuw et al., 2013; Zhao et al., 2001). However, other data showed that adding a repression or an activation domain to UFO changes its activity and that UFO is recruited at the *AP3* promoter in a LFY-dependent manner, rather suggesting a more direct role of UFO in gene regulation (Chae et al., 2008; Risseeuw et al., 2013). Hence, the molecular mechanism underlying LFY-UFO synergistic action remained elusive.

Here, we show that UFO connection to the SCF complex is partially dispensable for its activity and that an important role of UFO is to form a transcriptional complex with LFY at genomic sites devoid of canonical LFY binding sites (LFYBS). Our study presents a unique mechanism by which an F-box protein acts as an integral part of a transcriptional complex.

## RESULTS

### UFO F-box domain is partially dispensable for its floral role

To decipher the molecular mechanism underlying the synergistic action of LFY and UFO, we used a Dual Luciferase Reporter Assay (DLRA) to analyze their ability to activate promoters when transiently expressed in Arabidopsis protoplasts. We found that *pAP3,* the best characterized LFY-UFO target (Hill et al., 1998; Lamb et al., 2002), is strongly activated when LFY (or LFY-VP16, a fusion of LFY with the VP16 activation domain; Parcy et al., 1998) is co-expressed with UFO (or UFO-VP16) but not by either effector alone (Figure 1A and 1E). We also tested the promoter of *RABBIT EARS* (*RBE*), a gene expressed in petal primordia in a UFO-dependent manner (Krizek et al., 2006). *pRBE,* like *pAP3,* is specifically activated by LFY-UFO and LFY-VP16-UFO (Figure 1B) and not by the different versions of LFY or UFO alone. We also analyzed *pAP1* (the first 600 bp upstream *AP1* start codon) and *pAG* (a fusion of *AG* second intron with a minimal *35S* promoter), two LFY targets regulated by LFY independently of UFO (Busch et al., 1999; Parcy et al., 1998; Wagner et al., 1999). In the transient assay, *pAP1* and *pAG* activations by LFY and LFY-VP16 were insensitive to the addition of UFO (Figure 1C and 1D). Thus, the protoplast assay accurately reproduces several mechanisms of floral promoter activations.

**Figure 1.**
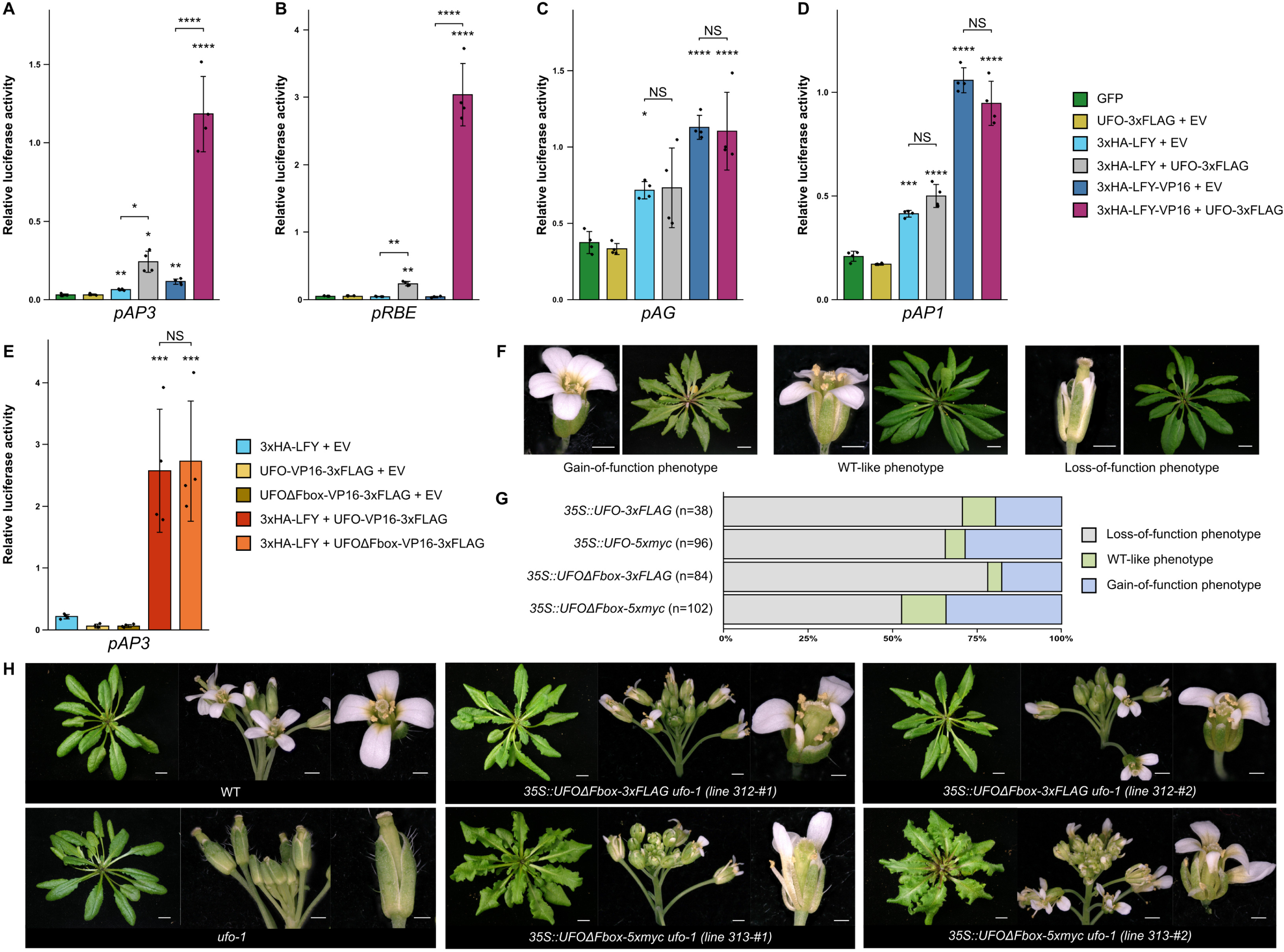
UFO action is largely independent on its F-box domain. (A-E) Promoter activations measured by DLRA. Effectors are indicated in the legend (right) and tested promoters below each graph. EV = Empty Vector. Data represent averages of independent biological replicates and are presented as mean ± SD, each dot representing one biological replicate (n = 4). One-way ANOVA with Tukey’s multiple comparisons test (C-D) or Welch’s ANOVA with Games-Howell post-hoc test (A, B and E). Stars above bars represent a significant statistical difference compared to the GFP control (A-D) or to 3xHA-LFY+EV (E), non-significant (NS) otherwise. (NS: p > 0.05,*: p < 0.05, **: p < 0.01; ***: p < 0.001 and ****: p < 0.0001). (F) Representative pictures of the different phenotypic classes obtained in the T1 population of WT plants overexpressing tagged versions of UFO or UFOΔFbox. Pictures are from different *35S::UFO-5xmyc* plants (scale bars, 1 mm for flowers and 1 cm for rosettes). (G) Distribution of T1 plants in phenotypic classes as described in (F). n = number of independent lines. Note that the severity of the phenotype varies within each class. (H) *ufo-1* complementation assay by the *35S::UFOΔFbox* transgene. Rosette (left, scale bar, 1 cm), inflorescence (middle, scale bar, 1 mm) and flower (right, scale bar, 0.5 mm) are shown. For each construct, two lines were crossed to *ufo-1* and at least 5 plants were analyzed per line. See also Figure S1F.

We next used this transient system to investigate the involvement of a SCF^UFO^-dependent ubiquitination pathway in *pAP3* activation by LFY-UFO. UFOdelF, a UFO version internally deleted of its F-box domain (and thus unable to insert into a SCF complex) was previously shown to be inactive, even inducing loss-of-function phenotypes when overexpressed in plants (Risseeuw et al., 2013). Such phenotypes were interpreted as a consequence of the inability of UFOdelF to ubiquitinate target proteins. To further investigate this point, we created a truncated UFO version (UFOΔFbox; aa. 91-443) in which the whole UFO N-terminal region, including the F-box domain, was deleted. In the protoplast assay, UFOΔFbox and UFOΔFbox-VP16, as opposed to UFOdelF, were able to activate *pAP3* when co-expressed with LFY (Figure 1E and S1A-C). Thus, the connection of UFO to an SCF complex appears dispensable for the transient *pAP3* activation. This suggests that UFOdelF is inactive *in planta* likely because its internal deletion affects its folding, and not because it lacks the connection to the SCF ubiquitination pathway.

We also stably expressed tagged versions of UFO and UFOΔFbox under the control of the constitutive *35S* promoter in Arabidopsis WT. In the T1 population of both types of transgenics (with or without the F-box), we observed phenotypes typical of *UFO* gain-of-function including serrated leaves, abnormal gynoecium, occasional extra-petals and stamens and reduced fertility (Figure 1F and 1G; Lee et al., 1997). Other T1 plants displayed reduced petal and stamen number resembling *ufo* mutants, but those showed no UFO protein expression (Figure S1D). Thus, both UFO and UFOΔFbox induce a gain-of-function phenotype in a WT background, in contrast to UFOdelF which induces a loss-of-function phenotype (Risseeuw et al., 2013).

To further evaluate the functionality of transgene-encoded tagged UFO and UFOΔFbox, we crossed gain-of-function transgenic lines to the strong *ufo-1* mutant (Wilkinson and Haughn, 1995). Both *35S::UFO* and *35S::UFOΔFbox* transgenes complemented *ufo-1* mutant and exhibited a gain-of-function phenotype (Figure 1H and S1E-F). Some defects such as missing or misshapen petals and disorganized flowers were specifically observed in the absence of the F-box, suggesting that this domain might be important for some UFO functions (Figure 1H and S1G). However, UFO and UFOΔFbox overall have a very similar activity, showing that the role of the F-box domain is largely dispensable. This suggested another molecular mechanism explaining the LFY-UFO synergy.

### The LFY-UFO complex binds a non-canonical sequence from *pAP3*

Protoplast assays established that *AP3* and *RBE* promoter sequences contain the information that dictates their specific activation by LFY-UFO. Thus, we searched for the *pAP3* element(s) required for such activation. Several regulatory regions driving *AP3* expression in early floral meristem have been identified, including the Distal Early Element (DEE) and the Proximal Early Element (PEE; Figure 2A; Hill et al., 1998). The DEE contains a predicted canonical LFYBS (Lamb et al., 2002; Figure S2A). Replacing this site by the high-affinity LFYBS from *pAP1* did not enhance activation (Figure S2B) and mutating it only slightly reduced *pAP3* activation (Figure S2C). Hence, in protoplasts like in plants (Lamb et al., 2002), *pAP3* LFYBS at DEE is not sufficient to explain *pAP3* activation. Previous studies suggested that LFY-UFO activation might involve the PEE and the region directly upstream (Hill et al., 1998; Lamb et al., 2002; Tilly et al., 1998). Indeed, in the protoplasts assay, we found that deleting the 107-bp region upstream of the PEE reduced LFY-UFO-dependent *pAP3* activation (Figure 2B). Refining the mapping, we identified a 20-bp region devoid of canonical LFYBS but important for LFY-UFO-dependent activation (Figure 2C and S2D).

**Figure 2.**
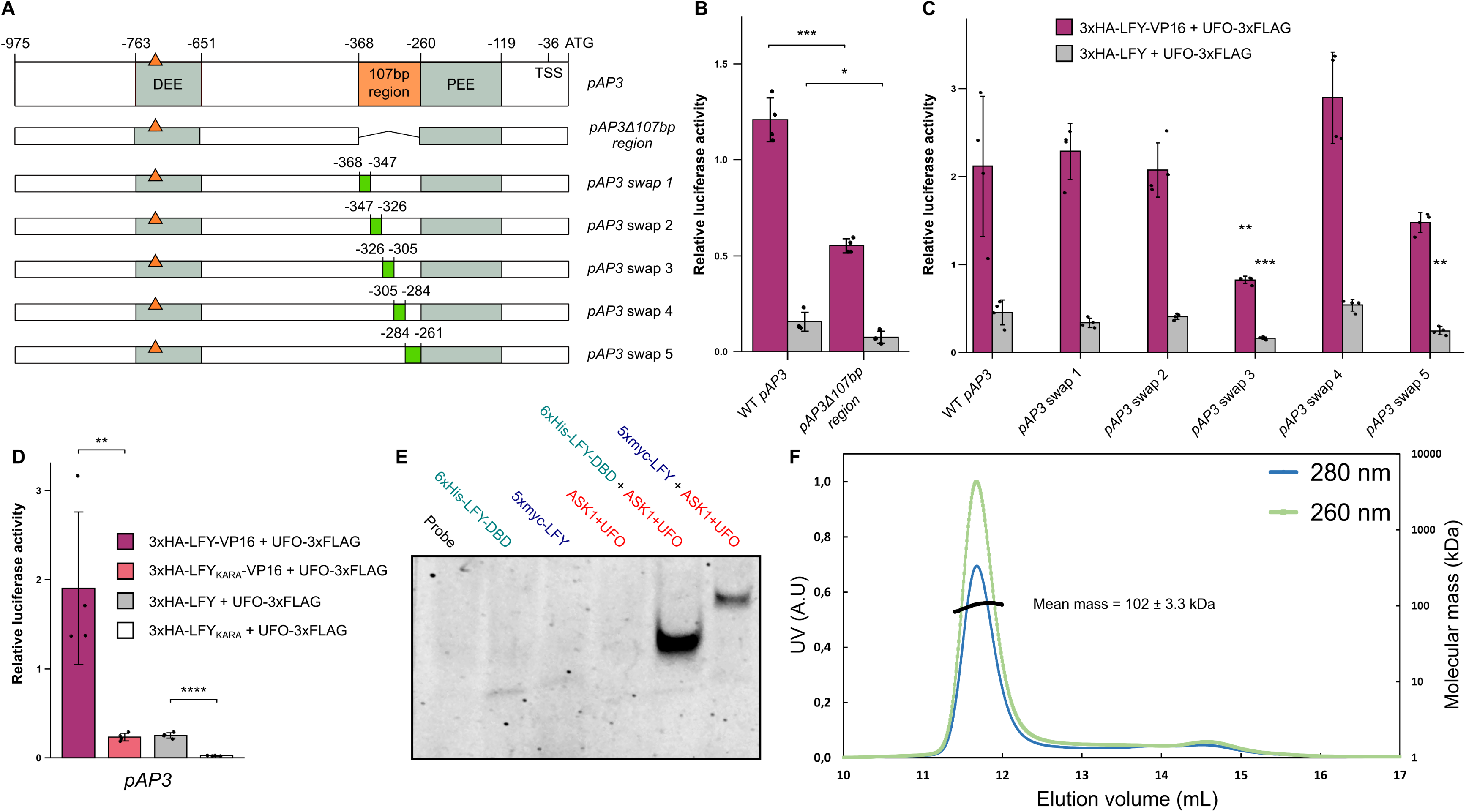
LFY and UFO form a transcriptional complex. (A) Description of *pAP3*. Top line represents WT *pAP3* with regulatory regions and *cis*-elements. Coordinates are relative to *AP3* start codon. TSS: Transcription Start Site. Orange triangle represents LFYBS. Other rows show the promoter versions used in (B-C). Green rectangles in swapped versions correspond to the same random sequence. (B and C) *pAP3* LFY-UFO response element mapping with *pAP3* versions described in (A) by DLRA in Arabidopsis protoplasts. See also Figure S2D. (D) Effect of the LFY KARA mutation (K303A-R233A) on *pAP3* activation in Arabidopsis protoplasts. See also Figure S2E and S2F. (E) EMSA with LUBS0 DNA probe and indicated proteins. Gel was cropped and only protein-DNA complexes are shown (see Supplemental Item 1). (F) Molecular mass determination for ASK1-UFO-LFY-DBD in complex with LUBS0 DNA by SEC-MALLS. Elution profiles correspond to absorbance at 280 nm and 260 nm (left ordinate axis, A.U: Arbitrary Unit). The black line shows the molecular mass distribution (right ordinate axis). See also Figure S1H. For bar charts, data represent averages of independent biological replicates and are presented as mean ± SD, each dot representing one biological replicate (n = 4). One-way ANOVA with Tukey’s multiple comparisons test (C). In (C), one-way ANOVA was performed with data from the same effector, and stars represent a statistical difference compared to WT *pAP3*. Unpaired t-tests (B and D). (NS: p > 0.05,*: p < 0.05, **: p < 0.01; ***: p < 0.001 and ****: p < 0.0001).

The absence of canonical LFYBS in *pAP3* elements required for LFY-UFO activation led us to examine whether LFY-DBD residues known to engage in direct contact with bases of the canonical LFYBS were required for *pAP3* activation. We found that mutations disrupting LFY binding on canonical LFYBS (LFY_K303A-R233A_ or LFY_KARA_; Chahtane et al., 2013) also strongly reduced *pAP3* activation even in the absence of the canonical LFYBS (Figure 2D and S2E). Importantly, this is not due to a compromised LFY-UFO interaction as LFY_KARA_ still interacts with UFO (Figure S2F). Thus, *pAP3* activation likely requires LFY DNA binding by some of the residues interacting with canonical LFYBS bases.

We investigated the possibility that LFY and UFO form a complex with a DNA element devoid of a canonical LFYBS by performing Electrophoretic Mobility Shift Assay (EMSA) with a DNA probe derived from the mapping experiment. Purified ASK1-UFO complex was combined either with recombinant LFY-DBD or with *in vitro-*produced Full Length (FL) LFY. None of the proteins (ASK1-UFO or LFY) bound the DNA probe alone but both LFY-DBD and FL LFY formed a complex with the DNA probe when mixed with ASK1-UFO (Figure 2E). Thus, a presumptive ASK1-UFO-LFY complex is formed on a *pAP3* DNA element (hereafter named LFY-UFO Binding Site 0 or LUBS0) that each partner does not bind on its own. However, when performing EMSA with low competitor DNA concentration, ASK1-UFO was able to bind the DNA probe, revealing a low affinity of ASK1-UFO for DNA (Figure S2G).

Using Size Exclusion Chromatography coupled to Multi-Angle Laser Light Scattering (SEC-MALLS), we determined a mass of 102 ± 3.3 kDa for this ASK1-UFO-LFY-DBD-LUBS0 complex, consistent with the presence of one copy of each protein per DNA molecule (theoretical mass of 108 kDa; Figure 2F and S2H). Mutating LUBS0 on various bases provided evidence that the formation of the complex is sequence-specific and suggested that the DNA motif might be bipartite (Figure S2I).

### Identification of the sequence motif bound by LFY-UFO

In order to identify all genome regions possibly targeted by the ASK1-UFO-LFY complex and obtain a precise definition for the LUBS motif, we performed ampDAP-seq (amplified DNA Affinity Purification sequencing; O’Malley et al., 2016). This technique allows the identification of binding sites of a recombinant protein or complex on naked genomic DNA (depleted of nucleosomes and methylation marks). We reconstituted the complex by mixing *in vitro*-produced 5xMyc-LFY and recombinant ASK1-UFO-3xFLAG complex with Arabidopsis genomic DNA, and sequenced and mapped the Myc-immunoprecipitated DNA fragments (Figure S3A). *pAP3* examination revealed two ASK1-UFO-LFY ampDAP-seq peaks, roughly located on the PEE and DEE, and similar to peaks obtained in LFY ChIP-seq experiments (Figure 3A; Goslin et al., 2017; Sayou et al., 2016). This contrasts with LFY ampDAP-seq (Lai et al., 2021) where a single peak was found, centered on the DEE canonical LFYBS. Similarly, *pRBE* contains a peak with ASK1-UFO-LFY in ampDAP-seq, but none with LFY alone (Figure S3B).

**Figure 3.**
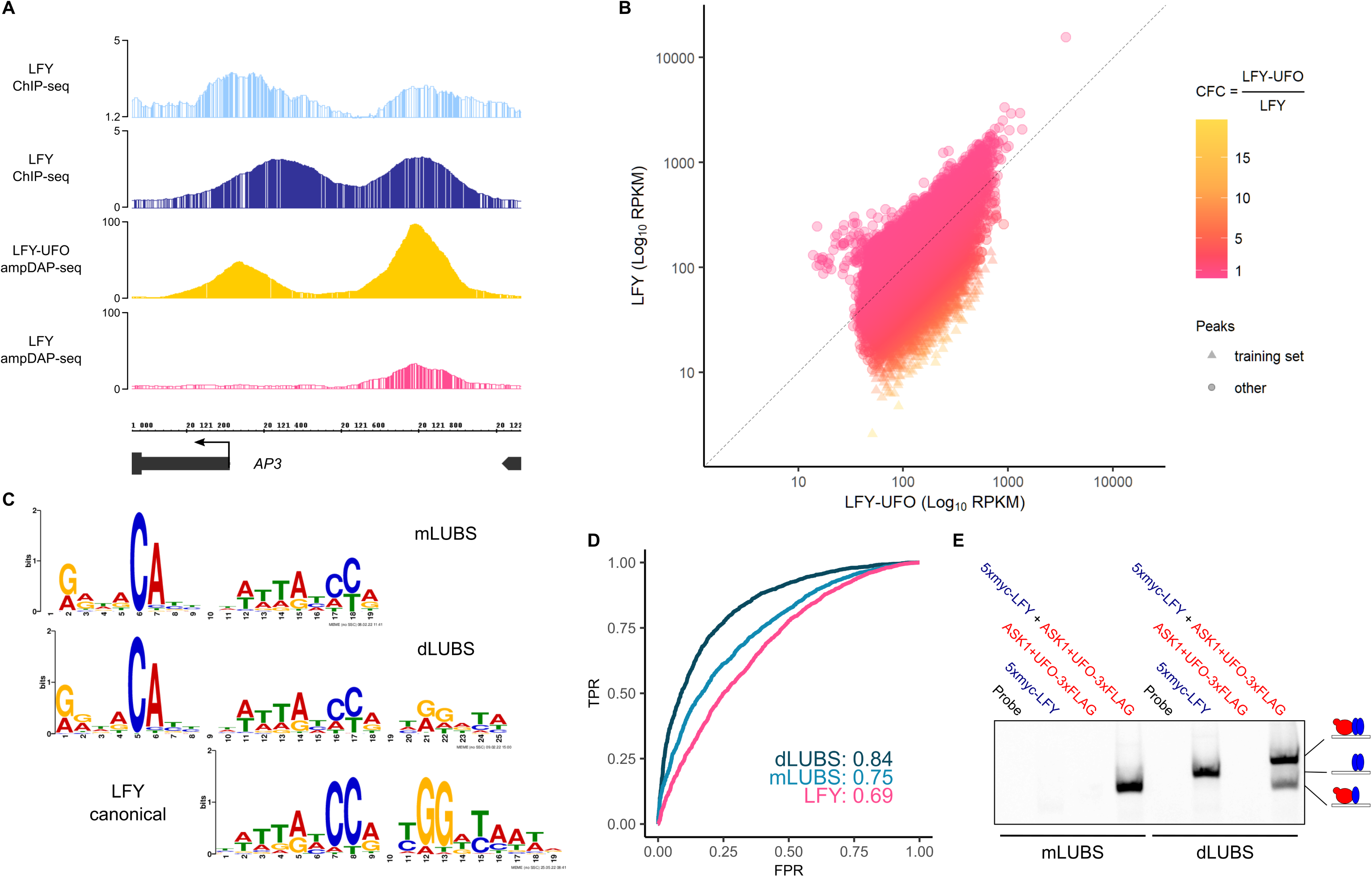
The LFY-UFO complex binds a novel DNA motif. (A) Integrated Genome Browser (IGB) view of *pAP3* showing LFY ChIP-seq in inflorescences (light blue; Goslin et al., 2017) or seedlings (dark blue; Sayou et al., 2016), LFY-UFO ampDAP-seq (yellow; this study), LFY ampDAP-seq (pink; Lai et al., 2021), y axis indicates read number range. (B) Comparison of peak coverage in LFY (y-axis) and LFY-UFO (x-axis) ampDAP-seq experiments, colored by CFC (peak coverage ratio in the presence or absence of UFO). LFY-UFO-specific peaks used to build mLUBS and dLUBS shown in (C) are triangle-shaped. (C) Logos for monomeric (mLUBS), dimeric (dLUBS) LFY-UFO binding sites and LFY binding site generated from ampDAP-seq experiments. The LFY logo was generated using the 600 peaks with the strongest LFY ampDAP-seq signal. (D) Receiver operating characteristics (ROC) curves for mLUBS, dLUBS and LFY using the top 20% high-CFC LFY-UFO-specific peaks. Area under the curve (AUC) values are shown. TPR: True Positive Rate, FPR: False Positive Rate. (E) EMSA with mLUBS and dLUBS highest score sequence DNA probes. Drawings represent the different complexes with LFY (blue) and ASK1-UFO (red) on DNA. Gel was cropped and only protein-DNA complexes are shown (see Supplemental Item 1). See also Figure S3F.

To find the LFY-UFO binding site, we selected regions where LFY binding highly depends on the presence of ASK1-UFO. For this, we computed the ratio between the coverage of peaks in the presence or absence of ASK1-UFO (this ratio was named Coverage Fold Change or CFC; Figure 3B). Searching for enriched DNA motifs in the 600 regions with the highest CFC (CFC > 4.7), we identified two bipartite motifs each made of two sequences separated by a variable region of fixed size. For both motifs, a 6-bp RRNRCA (N=A/C/G/T, R=A/G) sequence with a high-information CA is found at the 5’ side and the 3’ sequence resembles either a monomeric or a dimeric canonical LFYBS (Figure 3C). These LFYBS motifs present lower information content, i.e. more variability at each position compared to the canonical LFYBS. We named these motifs mLUBS and dLUBS for monomeric and dimeric LFY-UFO Binding Sites, respectively (Figure 3C). Since it is observed specifically when ASK1-UFO is present with LFY, the RRNRCA element will be next referred to as a UFO Recruiting Motif (URM). We modeled m- and d-LUBS using Position Weight Matrices (PWM), and tested their capacity to predict binding to the LFY-UFO-specific, high-CFC regions (top 20%). dLUBS PWM (and to a lesser extent mLUBS) outperformed LFY canonical PWM, showing that it better captured the ASK1-UFO-LFY specificity (Figure 3D). The LFYBS present within the LUBS of high CFC regions tended to have a lower PWM score (and thus lower predicted affinity) than LFYBS present in regions bound by LFY alone (Figure S3D), explaining why LFY binding to those sequences occurs only with UFO and the URM sequence. Remarkably, the URM was also identified *de novo* from published LFY ChIP-seq data (Goslin et al., 2017) by searching for enriched DNA motifs present at fixed distances from canonical LFYBS in regions bound by LFY *in vivo* but not by LFY alone *in vitro* (Figure S3E).

We validated ampDAP-seq findings by EMSA mixing DNA probes corresponding to optimal mLUBS and dLUBS motifs (highest score sequences for URM and LFYBS) with ASK1-UFO-LFY (Figure 3E and S3F). We observed a complex of slower mobility with dLUBS as compared to mLUBS, consistent with the presence of two LFY molecules on dLUBS. We also found that ASK1-UFO is able to supershift both FL LFY and LFY-DBD bound to canonical LFYBS from *pAP1* and *pAP3* DEE (Figure S3G), sometimes (but not systematically) increasing apparent LFY binding.

### LUBS are functional regulatory elements *in planta*

Next, we investigated the functional importance of LUBS *in vivo*. Using m- and d-LUBS PWMs, a medium score LUBS was identified in *pRBE* sequence (Figure 4A), bound by ASK1-UFO-LFY in EMSA (Figure S4A). Mutating either the URM, that abolished complex binding *in vitro* (Figure S4A), or the whole LUBS strongly reduced *pRBE* activation in protoplasts (Figure 4B). The functional importance of *pRBE* LUBS was also tested in Arabidopsis plants, where this promoter drives proper *RBE* expression in petal primordia (Takeda et al., 2004). Whereas the expected *RBE* expression pattern was observed in about half of our *pRBE::GUS* reporter plants, it was never found in plants with a mutated LUBS (Figure 4C and S4B), establishing its importance for the *RBE* activation.

**Figure 4.**
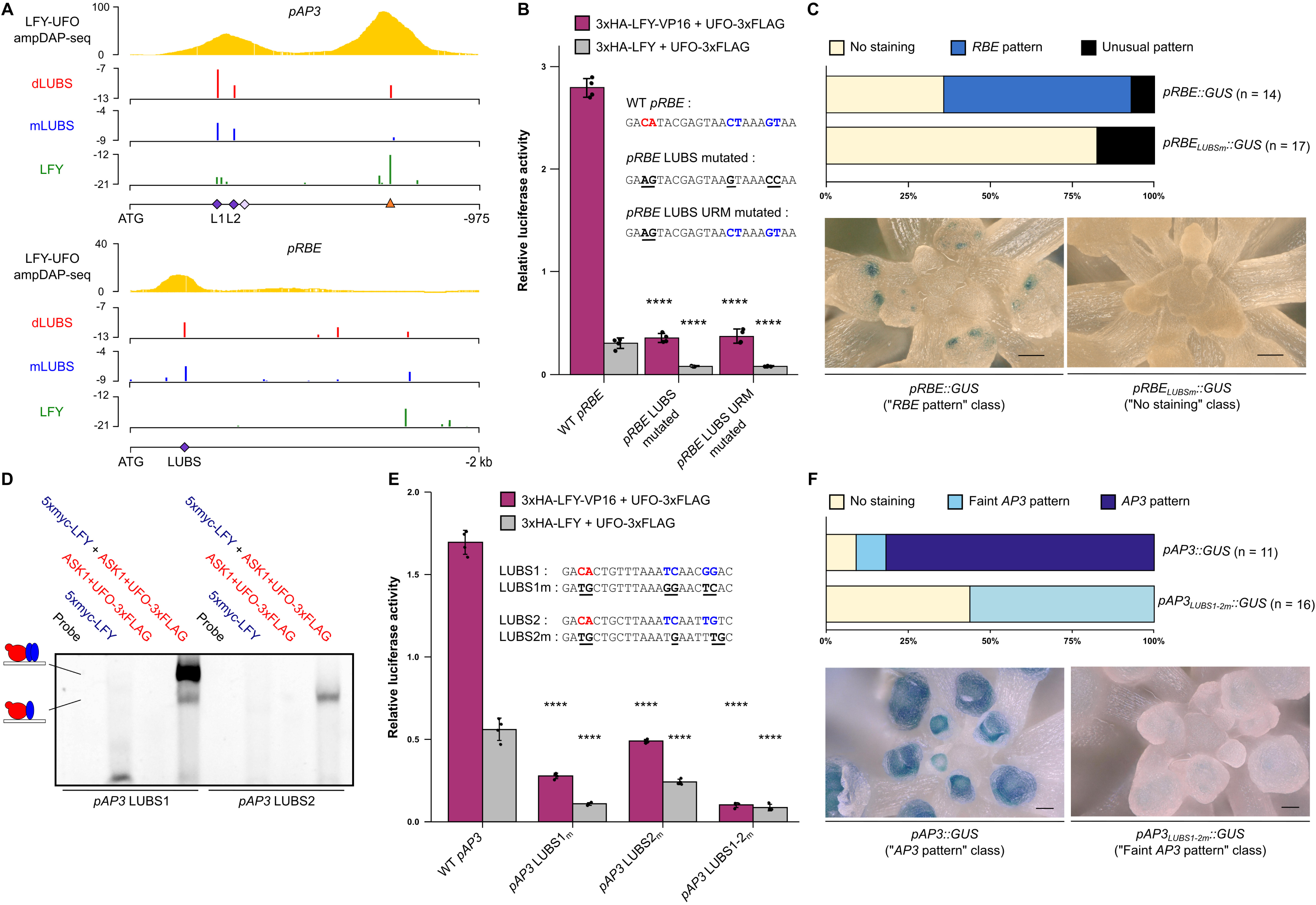
LUBS are required for LFY-UFO-dependent activations. (A) Identification of LUBS in *pAP3* and *pRBE*. IGB view of LFY-UFO ampDAP-seq with y-axis indicating read number range (top). Predicted binding sites using dLUBS and mLUBS models from Figure 3C and LFY PWM with dependencies (Moyroud et al., 2011), y-axis represents score values (bottom). The best binding sites correspond to the less negative score values. Studied LUBS, notably *pAP3* LUBS1 and LUBS2 (L1 and L2) are indicated with purple squares, *pAP3* canonical LFYBS as an orange triangle. LUBS0 (light purple square) is not visible because of its low score (mLUBS= -10.8; dLUBS= -25.6). (B) *pRBE* activation in Arabidopsis protoplasts. Effect of mutations (underlined) in URM (red) and in LFYBS (blue) bases of *pRBE* LUBS were assayed. (C) *In vivo* analysis of *pRBE::GUS* fusions. The percentage of transgenic lines with *RBE* pattern, unusual pattern or absence of staining was scored (top). n = number of independent lines. Unusual pattern refers to staining in unexpected tissues, each pattern seen in a single line. Representative pictures of plants with no staining (bottom left) and a *RBE* pattern (bottom right) are shown (scale bar, 50 µm). See also Figure S4B. (D) EMSA with *pAP3* DNA probes. Drawings represent the different complexes involving LFY (blue) and ASK1-UFO (red) on DNA. Gel was cropped and only protein-DNA complexes are shown (see Supplemental Item 1). (E) *pAP3* activation in Arabidopsis protoplasts. Effect of mutations (underlined) in URM (red) and LFYBS (blue) bases of *pAP3* LUBS were assayed. See also Figure S4G. (F) *In vivo* analysis of *pAP3::GUS* fusions. The percentage of transgenic lines with an *AP3* pattern, a faint *AP3* pattern or absence of staining was scored (top). n = number of independent lines. Faint *AP3* pattern corresponds to lines in which staining was observed in the expected domain but with a highly reduced intensity compared to normal *AP3* pattern. Representative pictures of plants with a faint *AP3* pattern (bottom left) and an *AP3* pattern (bottom right) are shown (scale bar, 50 µm). See also Figure S4H. For bar charts, data represent averages of independent biological replicates and are presented as mean ± SD, each dot representing one biological replicate (n = 4). One-way ANOVA with Tukey’s multiple comparisons test (B) or Welch’s ANOVA with Games-Howell post-hoc test (E). One-way ANOVA were performed with data from the same effector, and stars represent a statistical difference compared to WT promoters. (NS: p > 0.05,*: p < 0.05, **: p < 0.01; ***: p < 0.001 and ****: p < 0.0001).

We also searched for LUBS in *pAP3* and to our surprise, we identified several predicted sites of better score than LUBS0 in the PEE area (Figure 4A). The two highest score sites, LUBS1 and LUBS2, are specifically bound in EMSA by LFY in the presence of ASK1-UFO but not by ASK1-UFO alone (Figure 4D). A similar binding is also observed when combining LFY and UFOΔFbox (Figure S4C and S4D), consistent with *pAP3* activation assays from Figure 1. In the protoplast assay, mutating LUBS1 or LUBS2 significantly reduced *pAP3* activation (Figure 4E) with a stronger effect of the LUBS1 mutation, consistent with its high affinity for ASK1-UFO-LFY. Combining mutations in LUBS1 and in LUBS2 resulted in a drastic reduction of *pAP3* activation. Specifically mutating the URM of *pAP3* LUBS1 and LUBS2, that abolished LFY-UFO binding on individual sites in EMSA (Figure S4E and S4F), also reduced *pAP3* activation, albeit less effectively than mutating the whole LUBS (Figure S4G). Finally, the impact of *pAP3* LUBS mutation was evaluated in stable Arabidopsis transgenic plants. The previously described *pAP3::GUS* staining pattern in the second and third whorls of early floral meristems was severely reduced when LUBS1 and LUBS2 were mutated (Figure 4F and S4H).

Our analyses thus established the functional importance of *pAP3* and *pRBE* LUBS. However, LFY and UFO perform other functions together (Hepworth et al., 2006; Levin and Meyerowitz, 1995), suggesting that they likely target other genes. To identify possible LFY-UFO targets, we established a list of genes fulfilling the following criteria. These genes should be i) present in the vicinity of LFY-UFO specific peaks in ampDAP-seq (high CFC) ii) bound *in vivo* in LFY ChIP-seq experiments (Goslin et al., 2017; Jin et al., 2021; Moyroud et al., 2011; Sayou et al., 2016) and iii) deregulated in *ufo* inflorescences (Schmid et al., 2005). This list (Figure S4I) includes *AP3* and the other B gene *PISTILLATA* (*PI*), previously proposed as a LFY-UFO target (Honma and Goto, 2000) but through an unknown regulatory element that our LUBS model precisely localized (Figure S4J). We also found floral regulators such as SQUAMOSA PROMOTER BINDING PROTEIN-LIKE 5 (SPL5) and FD as well as novel candidates involved in cell wall synthesis or cytokinin catabolism, two processes important for floral meristem emergence (Figure S4I).

### The LFY K249R mutation specifically affects UFO-dependent LFY functions

Next, we wondered whether UFO-dependent and independent LFY functions could be decoupled. We took advantage of *pAP3* activation in protoplasts to search for LFY mutations specifically impairing LFY-UFO synergistic action. As we initially looked for ubiquitination mutants, we mutated exposed lysines from LFY-DBD into arginines (Hamès et al., 2008). Among three tested residues, the LFY K249R mutation (Figure S5A) strongly reduced *pAP3* activation by LFY-UFO (Figure 5A) or LFY-VP16-UFO (Figure S5B) but did not prevent the UFO-independent *pAG* activation (Figure S5C). Yeast-two-hybrid (Y2H) experiment showed that the LFY K249R mutation did not disrupt LFY-UFO interaction (Figure S5D) suggesting that it rather specifically affected the ASK1-UFO-LFY-DNA complex formation (Figure S5E). We tested this hypothesis genome-wide in ampDAP-seq (Figure S5F and S5G). We found that LFY_K249R_ alone binds canonical LFYBS as well as LFY (Figure S5H). However, the UFO-dependent LFY binding was strongly reduced by the K249R mutation (Figure 5B, 5C and S5I), showing that LFY K249 plays a key role in LUBS binding.

**Figure 5.**
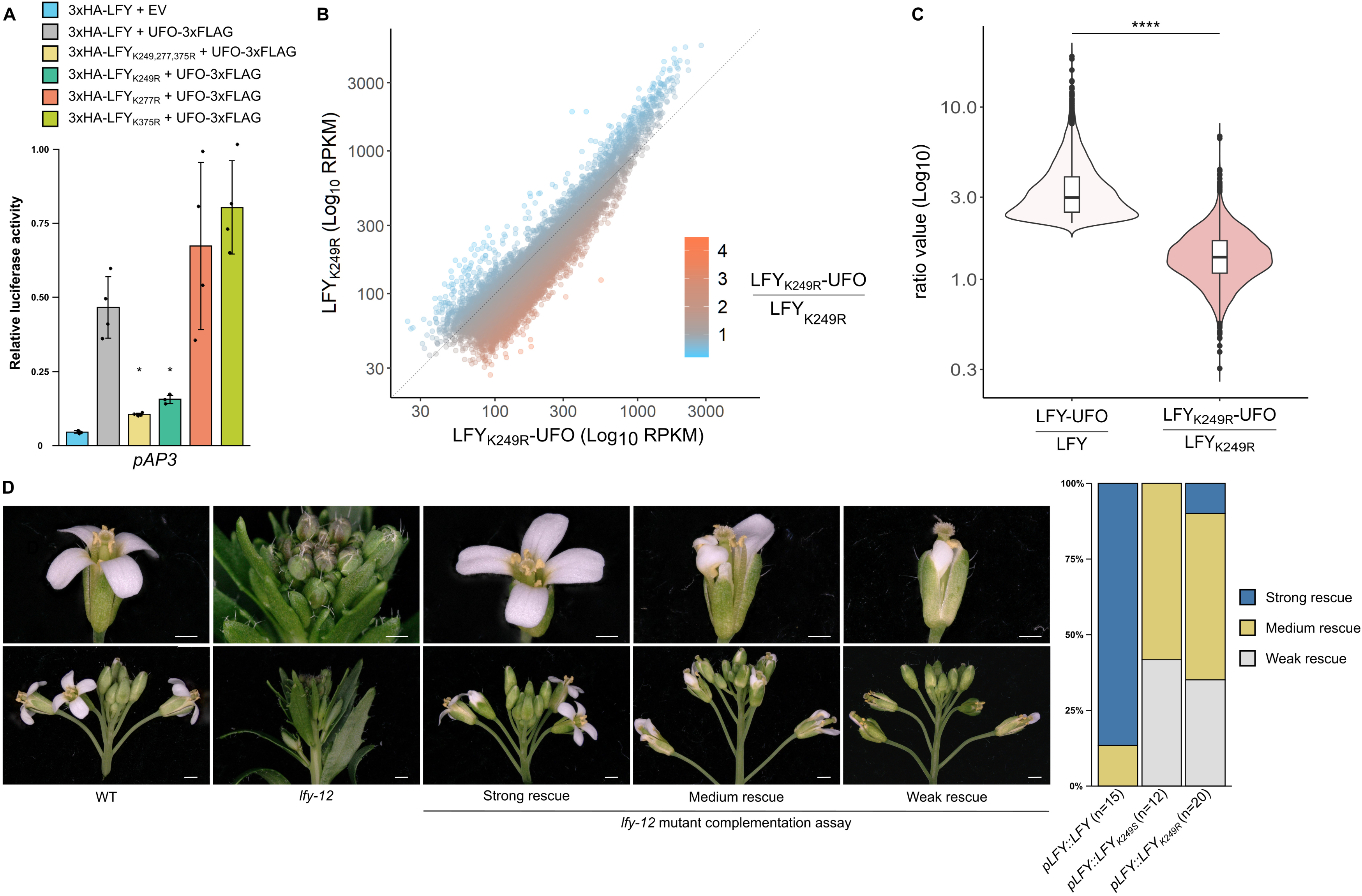
The LFY K249R mutation specifically disrupts the LFY-UFO synergy. (A) *pAP3* activation in Arabidopsis protoplasts. Data represent averages of independent biological replicates and are shown as mean ± SD, each dot representing one biological replicate (n = 4). Welch’s ANOVA with Games-Howell post-hoc test. Stars indicate a statistical difference compared to 3xHA-LFY+UFO-3xFLAG. (NS: p > 0.05,*: p < 0.05). See also Figure S5B and S5C. (B) Comparison of peak coverage in LFY_K249R_ (y-axis) and LFY_K249R_-UFO (x-axis) ampDAP-seq experiments, colored by peak coverage ratio. Note that, in contrast with Figure 3B, UFO weakly affects LFY_K249R_ binding genome-wide. (C) Effect of LFY K249R mutation on LFY-UFO genome-wide binding. Violin plots show the distribution of coverage ratios for LFY and LFY_K249R_ for LFY-UFO-specific regions (20% highest CFC). Wilcoxon’s rank sum test (***: p < 0.0001). (D) *lfy-12* mutant complementation assay. Pictures of WT, *lfy-12* mutant and of representative plants of the different phenotypic complementation classes (left, scale bar 1 mm for the top and 1 cm for the bottom). Strong rescue picture was taken from a line expressing *pLFY::LFY*, medium and weak rescues from lines expressing *pLFY::LFY_K249R_*. Distribution of *pLFY::LFY*, *pLFY::LFY_K249S_* and *pLFY::LFY_K249R_* lines within phenotypic complementation classes (right). n = number of independent lines.

The importance of LFY K249 was tested in Arabidopsis plants using complementation assay of the *lfy-12* null mutant (Weigel et al., 1992). *lfy-12* plants expressing LFY_K249R_ or LFY_K249S_ under the control of *LFY* promoter developed flowers with normal sepals and carpels but with defective third and more importantly second whorl organs, resulting in flowers of weak *ufo* mutants (Figure 5D; Durfee et al., 2003). When expressed under the constitutive *35S* promoter, LFY_K249R_ triggered ectopic flower formation and early flowering like WT LFY (Figure S5J; Weigel and Nilsson, 1995), consistent with these LFY functions being independent of UFO and thus not affected by the K249R mutation.

### Structural characterization of the ASK1-UFO-LFY-DNA complex

The results we obtained so far demonstrate the existence and the functional importance of the LFY-UFO-LUBS complex but do not explain how the complex is able to recognize a novel DNA element. In particular, whether UFO interacts with LUBS DNA directly or instead changes LFY conformation triggering a novel contact between LFY-DBD (for example involving the K249 residue) and DNA is unknown. To gain insight into the properties of this newly identified complex, we reconstituted an ASK1-UFO-LFY-DBD-LUBS1 complex and we examined it using cryo-electron microscopy (Figure 6A and S6A-D). We obtained a structure of the complex at a medium resolution (4.27 Å; Figure S6G-I). We then fitted the AlphaFold2 predicted structure for UFO (Q39090), ASK1 (Q39255) and the LFY-DBD dimer/DNA crystallographic structure (PDB entry 2VY1; Hamès et al., 2008 ; Figure S6E) into the cryo-electron microscopy density map (Figure 6B and S6F). UFO, LFY-DBD, ASK1 and the DNA were clearly recognizable. In these experimental conditions, a mixture of two complexes was observed with either one or two LFY-DBD molecules (Figure S6C-D). As expected, ASK1 interacts with the F-box domain of UFO (Figure 6D).

**Figure 6.**
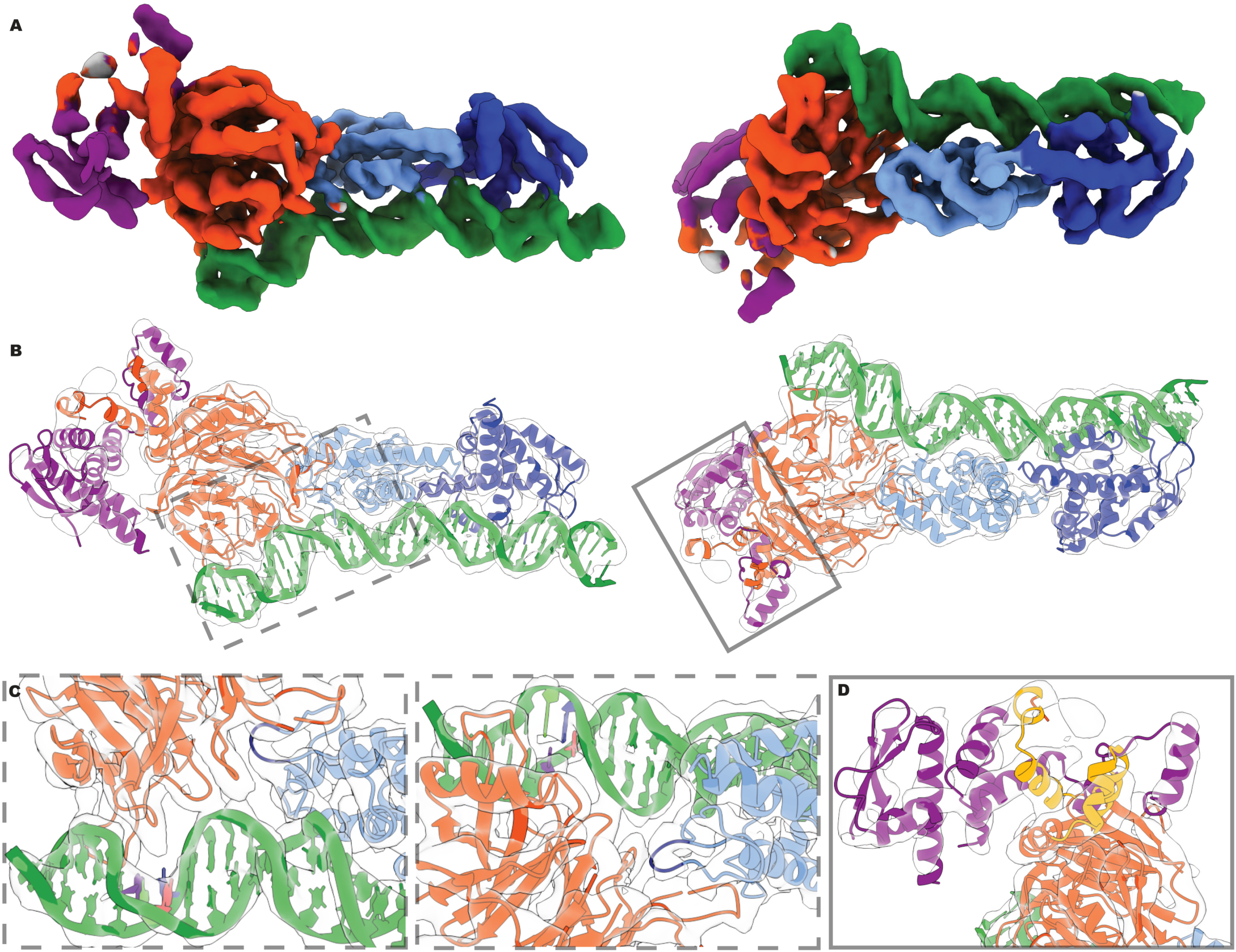
Structural characterization of the ASK1-UFO-LFY-DNA complex. (A) Cryo-EM density map of the ASK1-UFO-LFY-DBD-LUBS1 complex under two angles, colored with regard to the underlying macromolecule (green: LUBS1 DNA; pale and dark blue: LFY-DBD; red: UFO; purple: ASK1). (B) The same views of the cryo-EM density map in transparent gray with fitted structures of LFY-DBD dimer (pale and dark blue), UFO (red), ASK (purple) and DNA (dark green and filled rings in red for A, blue for T, pale green for G and yellow for C). The frames roughly indicate the regions shown in panels C (dashed line, two views) or D (filled line). (C) Zoom on the UFO-DNA contact region (left) and on the LFY-UFO interface (right). For clarity, only the high-information CA of the URM and its complement is highlighted by filled coloring the rings for each base. The loop of LFY-DBD that contains the K249 residue is highlighted in dark blue. (D) Zoom on the ASK1-UFO interface, with the F-box of UFO highlighted in gold.

The fitting revealed that at least four basic loops projecting from the UFO β-propeller makes direct contacts with the DNA major groove (around the URM area; Figure 6C), resulting in a bend of roughly 30° degrees in the DNA double helix (Figure S6F). The presence of these basic loops is consistent with the low affinity binding of ASK1-UFO to DNA (Figure S2G). Consistent with previous high-resolution crystal structures of LFY-DBD in complex with DNA (Hamès et al., 2008), LFY interacts with the LUBS via its helix-turn-helix motif, with the C-terminal helix of the HTH lying in the major groove and an N-terminal extension mediating additional contacts in the minor groove.

The complex structure also revealed an interface between UFO and one LFY-DBD monomer (Figure 6C). The LFY-DBD loop that bears the K249 residue lies in this interface and likely interacts with one of the DNA-binding loops of UFO, consistent with the key role of LFY K249 in the ternary complex formation. These data show how a β-propeller protein is able to modify the specificity of a TF, and explains how the LFY-UFO complex synergistically recognizes a novel DNA element.

### The LFY-UFO complex likely has a deep evolutionary origin

As genetic and physical LFY-UFO interactions have been described in diverse angiosperms, we wondered whether the mechanism unraveled for Arabidopsis proteins could also apply to LFY from other species, including non-angiosperm ones. We selected LFY orthologous proteins from several species and with different DNA binding specificities (Figure 7A). Indeed, through evolution, LFY specificity evolved with three major DNA binding specificities: type I (Figure 3C) in angiosperms, gymnosperms, ferns and in the moss *Marchantia polymorpha*, type II in the moss *Physcomitrium patens* and type III in algae (Sayou et al., 2014). Because functional UFO homologs have not been identified in all these plant groups, we used Arabidopsis UFO in all the following experiments.

**Figure 7.**
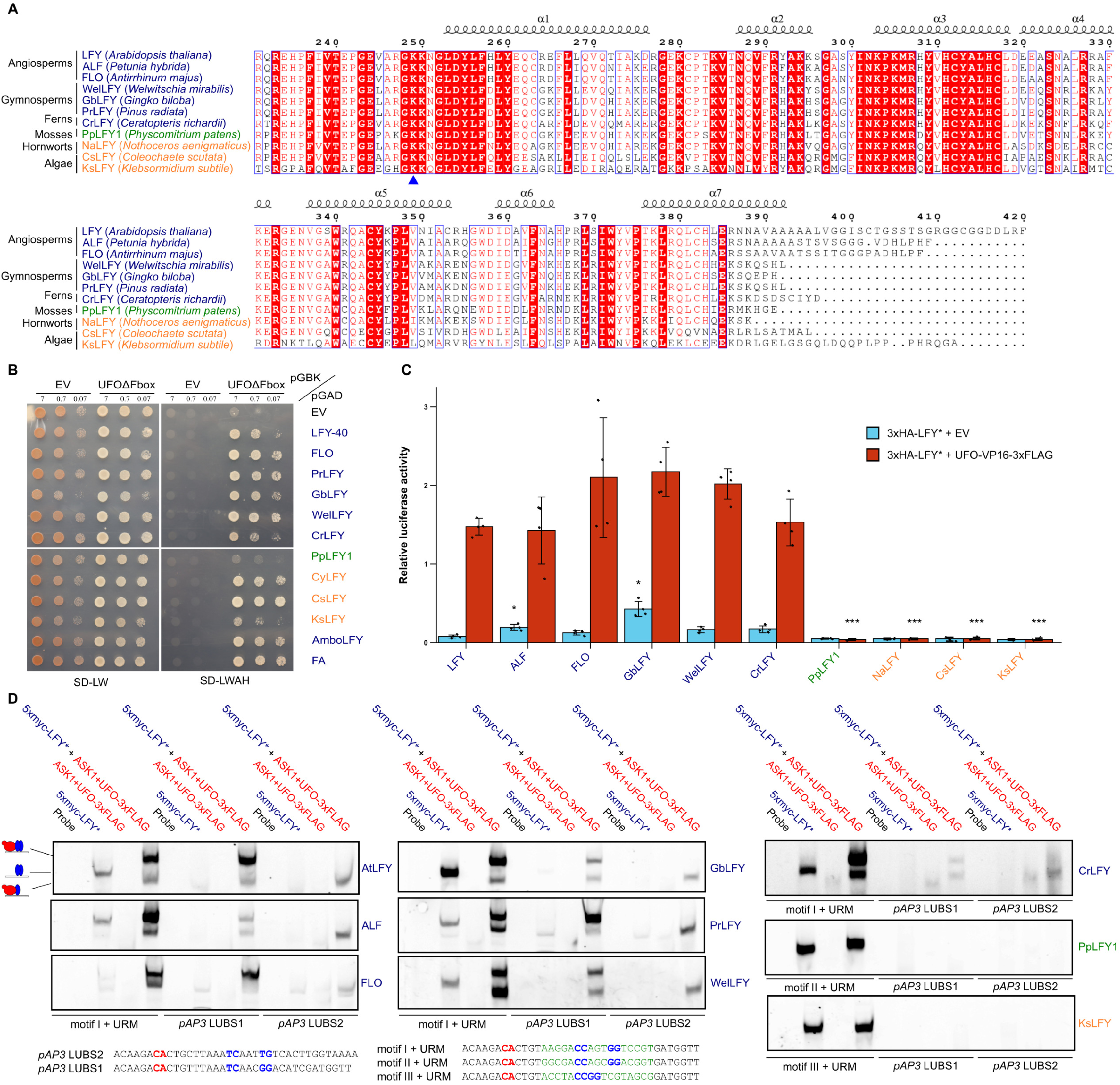
LFY-UFO interaction is conserved beyond angiosperm species. (A) Alignment of LFY DBDs. Amino acid numbering and secondary structure annotation are based on LFY from *A. thaliana*. LFY K249 residue is indicated with a blue triangle. DNA binding specificities are color-coded, type I (blue), II (green) and III (orange). FLO= FLORICAULA; ALF = ABERRANT LEAF AND FLOWER. (B) Interaction between LFY orthologs and AtUFOΔFbox in Y2H. LFY orthologs are described in (A) except CyLFY (*Cylindrocystis sp.*), AmboLFY (*Amborella trichopoda*) and FA (FALSIFLORA; *Solanum lycopersicum*). See Figure S2F for legends. (C) *pAP3* activation measured by DLRA in Arabidopsis protoplasts. EV = Empty Vector. 3xHA-LFY* refers to the different LFY orthologs indicated under the x-axis. Data represent averages of independent biological replicates and are presented as mean ± SD, each dot representing one biological replicate (n = 4). Welch’s ANOVA with Games-Howell post-hoc test. One-way ANOVA was performed with data from the same effector (described in the legend), and stars represent a statistical difference compared to AtLFY, NS otherwise. (NS: p > 0.05,*: p < 0.05, **: p < 0.01; ***: p < 0.001). (D) EMSA with indicated DNA probes (bottom). URM and LFYBS bases are depicted in red and blue, respectively. *pAP3* LUBS1 sequence was modified to insert the perfect sequence of motif I, II or III (depicted in green; Sayou et al., 2014): these DNA probes were used as positive controls for binding of LFYs alone and LFY-UFO complex formation. 5xmyc-LFY* refers to the different LFY orthologs indicated next to each EMSA and described in (A). Gels were cropped and only protein-DNA complexes are shown (see Supplemental Item 1).

We tested the interaction of various LFY orthologs with AtUFO in Y2H (Figure 7B), in DLRA in protoplasts with Arabidopsis *pAP3* (Figure 7C) and in EMSA (Figure 7D). In Y2H, all LFYs except LFY from *P. patens* (Type II) interact with AtUFO (Figure 7A). However only Type I LFY from angiosperms, gymnosperms and ferns form a complex on *pAP3* LUBS and activate *pAP3* in the protoplast assay. These results suggest that the ability of LFY and UFO to act together by forming a complex is ancient, largely predating the origin of angiosperms. We obtained no evidence that type II and III LFY (from moss and algae) could form a complex with AtUFO on LUBS1 and 2. A detailed history of the LFY-UFO interaction will await further analyses, notably with the identification of UFO orthologs from non-angiosperm genomes.

## DISCUSSION

LFY was long known to interact with UFO to induce the localized expression of the B floral homeotic gene *AP3* (and *PI,* albeit to a lesser extent) but the molecular nature of their synergistic action had remained unknown. As UFO encodes an F-box protein and belongs to an SCF complex (Ni et al., 2004; Wang et al., 2003; Zhao et al., 1999, 2001), it was thought to target proteins for a SCF^UFO^-dependent ubiquitination and possible degradation. LFY was an obvious target candidate but clear evidence of LFY ubiquitination was missing (Chae et al., 2008). The results we present here suggest that the F-box domain, required for ubiquitination, is dispensable for most UFO-dependent LFY activity. Nevertheless, the high conservation level of UFO F-box sequence in angiosperms, together with slight differences in UFO activity when the F-box is deleted suggest that this domain might still be needed for some facets of UFO function. It is for instance possible that UFO works redundantly with other F-box proteins in ubiquitination pathways, like with the F-box protein HAWAIIN SKIRT identified in a genetic screen as an enhancer of *ufo* mutant phenotype (Levin et al., 1998).

Our key finding is that UFO interacts with LFY and DNA to form a transcriptional complex at novel regulatory sites in the promoters of several regulators of floral organ development (including *AP3*, *PI* and *RBE*). These regulatory sites (mLUBS and dLUBS) are made of a low-affinity or half LFYBS (poorly or not bound by LFY alone) and a URM located at a fixed distance from it and responsible for UFO recruitment. The formation of such a sequence-specific complex is explained at the structural level by the capacity of UFO to interact with both LFY and DNA, independently of ubiquitination. The poor ability of ASK1-UFO to bind DNA alone explains its complete dependence towards LFY to interact with regulatory sites and perform its functions *in planta* (Lee et al., 1997; Risseeuw et al., 2013). Thus, depending on *cis*-elements, LFY either binds DNA as a homodimer or requires UFO to form a ternary complex. Mutation of the LFY K249 residue allows uncoupling these two types of binding by specifically disrupting the formation of the LFY-UFO-DNA complex. The position of this residue in the 3D structure at the interface between LFY, UFO and DNA is consistent with the key role of this residue in the complex formation. Obtaining a higher-resolution structure will help to understand precisely its interactions.

Very few other cases where non-TF proteins are recruited by a DNA motif at a fixed distance of a well-characterized TF have been described so far: it is the case for Met4 and Met28 with the TF Cbf1 in yeast (Siggers et al., 2011), or the herpes simplex virus transcriptional activator VP16 with the Oct-1/HCF-1 complex (Babb et al., 2001). None of these examples involves an F-box protein or a Kelch-type β-propeller protein and neither of them has been characterized at the structural level. Plant genomes have hundreds of Kelch-containing proteins and others may perform a similar function. More generally, it becomes increasingly clear that genome-wide binding of TFs *in vivo* is not entirely explained by direct binding by homo-complexes, as evidenced by the comparison between ChIP-seq and (amp)DAP-seq data (Lai et al., 2021; O’Malley et al., 2016). The mechanism by which cofactors (that do not necessarily bind DNA on their own) bring TF to specific loci might be common, and it is a broad field to explore in the future.

The molecular mechanism we discovered here is consistent with most published data on *AP3* and *PI* regulation (Chae et al., 2008; Hill et al., 1998; Honma and Goto, 2000; Tilly et al., 1998). However, precise understanding of regulation of *AP3* and *RBE* will require further work on other *cis* and *trans*-elements. For example, why *AP3* is not transcribed before floral stage 2-3 despite LFY and UFO expression, why *pAP3* is not activated by LFY (or LFY-VP16) alone through the canonical LFYBS or how *SEPALLATA3* constitutive expression allows *AP3* induction independently of UFO (Castillejo et al., 2005) are open questions.

In Arabidopsis, LFY and UFO expression also overlap in the earliest floral stages and likely regulate other genes to promote floral meristem identity or repress bracts (Hepworth et al., 2006). Moreover, in many other species, the joint action of LFY and UFO is more important than in Arabidopsis with a key role on the flower meristem fate in petunia and tomato or in panicle architecture in rice (Ikeda-Kawakatsu et al., 2012; Lippman et al., 2008; Souer et al., 2008). It is thus likely that the mechanism we discovered here will expand the molecular understanding of flower development in multiple species and for multiple targets genes. Since our data suggest that LFY interacts with UFO in non-angiosperm species, the LFY-UFO complex might also regulate the development of reproductive cones in gymnosperms or lateral meristems in ferns (Moyroud et al., 2017; Plackett et al., 2018).

## Limitation of this study

The UFO ubiquitination-independent role was evidenced by the fact that UFOΔFbox rescues many *ufo-1* defects when overexpressed in Arabidopsis, but this experiment was not performed with *UFO* endogenous promoter. It is possible that, when UFO is expressed at wild type levels, the need for ubiquitination becomes more obvious. Whether the connection to the ubiquitination pathway is required with the LFY-UFO-DNA complex or within another molecular context involving UFO and another substrate in unknown.

The role of ASK1 was not studied in detail here, but we noticed in some biochemical experiments that ASK1 seems to facilitate the formation of the ASK1-UFO-LFY-DNA complex *in vitro.* It can be hypothesized that ASK1 not only has a role in ubiquitination but also helps to stabilize the LFY-UFO complex on DNA *in vivo*.

Finally, the low resolution of the structure precluded to precisely position the UFO protein and to identify the residues and the chemical bonds involved in the UFO-DNA and UFO-LFY interfaces.

## ACKNOWLEDGMENTS

We thank AM. Boisson for preparing suspension cells, X. Lai for ampDAP-seq libraries and technical assistance and R. Koes for sharing data and materials. We gratefully acknowledge C. Marondedze, G. Vachon, M. Le Masson, C. Berthollet, B. Orlando Marchesano and J. Bourenane-Vieira for help with experiments. We thank G. Vert, U. Dolde and R. Dumas for discussion. The electron microscopy facility is supported by the Rhône-Alpes Region, the FRM, the FEDER and the GIS-IBISA. This work used the platforms of the Grenoble Instruct-ERIC center (ISBG; UAR 3518 CNRS-CEA-UGA-EMBL) within the Grenoble Partnership for Structural Biology (PSB), supported by FRISBI (ANR-10-INBS-0005-02). We thank Caroline Mas for assistance and access to the biophysical platform. This work was supported by the GRAL Labex financed within the University Grenoble Alpes graduate school (Ecoles Universitaires de Recherche) CBH-EUR-GS (ANR-17-EURE-0003), the CEA (PhD fellowship to PR) and the ANR-17-CE20-0014-01 Ubiflor project to FP.

## AUTHOR CONTRIBUTIONS

FP and PR designed the project

PR performed the plant experiments helped by GT

PR and ET performed the biochemical experiments helped by HC for evolutionary analyses LT performed the bioinformatics analyses helped by JL and RBM

EZ and GS performed cryo-EM experiments and MN, EZ, CZ and GS analyzed the data PR and LT assembled the figures

PR and FP wrote the paper with contributions from LT and CZ

## STAR METHODS

### KEY RESOURCE TABLE

Provided as a separate file

### RESOURCE AVAILABILITY

#### Lead contact

Further information and requests for resources and reagents should be directed to and will be fulfilled by the Lead Contact, François Parcy.

#### Materials availability

Plasmids and transgenic lines generated in this study are available from the Lead Contact without restriction.

#### Data and code availability

ampDAP-seq data have been deposited at GEO and are publicly available as of the date of publication (GSE204793). All original code has been deposited at github (https://github.com/Bioinfo-LPCV-RDF/LFYUFO_project) and is publicly available as of the date of publication. Any additional information required to reanalyze the data reported in this paper is available from the lead contact upon request. The cryo-EM structure determined in this study is deposited in the EM data bank under the reference number EMD-15145.

### EXPERIMENTAL MODEL AND SUBJECT DETAILS

#### Arabidopsis growth

All mutants and transgenic lines are in the *A. thaliana* Columbia-0 accession. Seeds were sown on soil, stratified 3 days at 4 °C, and then grown at 22°C under long-day conditions (16 h light). Transgenic plants were obtained with *Agrobacterium tumefaciens* C58C1 pMP90 using the floral dip method. Transformants were identified using GFP or Basta selection.

#### Arabidopsis cell suspension culture

*Arabidopsis thaliana* (ecotype Columbia-0) cells in suspension cultures were grown under continuous light (90 μmol of photons m^-2^ s^-1^) at 21 °C with shaking at 135 rpm in Murashige and Skoog (MS) medium supplemented with 30 g/L sucrose and 2 mg/L 2,4-dichlorophenoxyacetic acid (2,4D), pH 5.5. Suspension cells were subcultured every week with a 5-fold dilution. Suspension cells at 4 or 5 days following subculture were used for protoplast preparation.

### METHOD DETAILS

#### Cloning

DNA fragments were amplified by PCR with Phusion high fidelity polymerase (NEB). Plasmids were all obtained by Gibson Assembly (GA) with either PCR-amplified or restriction enzyme-digested backbone vectors. We used the 420 aa LFY version. For site-directed mutagenesis, primers containing the desired mutations were used for GA mutagenesis. Plasmids were obtained using DH5α bacteria and were all verified by Sanger sequencing. A list of plasmids and cloning procedures is provided in Supplemental Table 1. Oligonucleotide sequences are listed in Supplemental Table 2.

#### Yeast-two-hybrid

Coding sequences were cloned in pGADT7-AD or pGBKT7 vectors (Clontech) by GA. Y187 and AH109 yeast strains (Clontech) were transformed with pGADT7-AD or pGBKT7 vectors and selected on plates lacking Leucine (SD-L) or Tryptophan (SD -W), respectively (MP Biomedicals). After mating, yeasts were restreaked on plates lacking Leucin and Tryptophan (SD -L-W) for 2 days. Yeasts were then resuspended in sterile water and OD_600nm_ was adjusted to indicated values for all constructions; two ten-fold dilutions were performed, and 6 μL drops were done on SD -L-W or SD -L-W-A-H (lacking leucine, tryptophan, histidine and adenine) plates. Yeasts were grown at 28 °C and pictures were taken at indicated times.

#### Dual Luciferase Reporter Assay in Arabidopsis protoplasts

Effector plasmids with a 3xHA tag were obtained by cloning indicated genes in the modified pRT104 vector containing a 3xHA N-terminal tag (pRT104-3xHA, Chahtane et al., 2018). The pRT104 empty plasmid (Topfer, 1987) was reengineered to insert a 3xFLAG C-terminal tag. For reporter plasmids, indicated promoter fragments were cloned upstream a Firefly Luciferase gene in pBB174 (Blanvillain et al., 2011). The pRLC reference plasmid contains Renilla Luciferase sequence under the control of the *35S* promoter. Plasmids were obtained in large amounts using NucleoBond Xtra Maxi Plus kit (Macherey-Nagel).

Protoplasts were prepared from Arabidopsis Col-0 cell suspension and transformed following the procedure described by Iwata et al. (Iwata et al., 2011). Cell wall was digested using Onuzuka R-10 cellulase and macerozyme R-10 (Yakult Pharmaceutical). Digested cells were passed through two layers of Miracloth to remove debris, and protoplast concentration was adjusted to 2-5x10^5^ cells/mL. Protoplasts were then PEG-mediated transformed using 10 μg of indicated effector and reporter plasmids and 2 μg of reference plasmid. After 17 h of incubation at RT, protoplasts were lysed. Firefly (F-LUC) and Renilla Luciferase (R-LUC) activities were measured using Dual Luciferase Reporter Assay System (Promega) and a TECAN Spark 10M 96-well plate reader. F-LUC/R-LUC luminescence ratios were calculated with background-corrected values. Four biological replicates were done for each plasmid combination.

#### Electrophoretic Mobility Shift Assay (EMSA)

DNA probes used in EMSA are listed in Supplemental Table 2. Complementary oligos were annealed overnight in annealing buffer (10 mM Tris pH 7.5, 150 mM NaCl and 1 mM EDTA). 4 pmol of double-stranded DNA was then fluorescently labeled with 1 unit of Klenow fragment polymerase (NEB) and 8 pmol Cy5-dCTP (Cytiva) in Klenow buffer during 1 h at 37 °C. Enzymatic reaction was stopped with a 10-min incubation at 65 °C.

Proteins used in EMSA were obtained by different methods (bacteria, insect cells or TnT). Recombinant proteins (6xHis-LFY-DBD, UFOΔFbox-3xFLAG) and recombinant complexes (ASK1-UFO, ASK1-UFO-3xFLAG) concentration was adjusted to 500 nM for all reactions. All the 5xmyc-tagged proteins were obtained *in vitro* by TnT. 50 µL TnT reactions were done by mixing for 2 h at 25 °C 5 µg of pTNT-5xmyc plasmid containing the gene of interest with TnT SP6 High-Yield Wheat Germ Protein Expression System (Promega). For EMSA with TnT-produced proteins, 5 µL of TnT reaction was used. Recombinant protein buffer or TnT mix was used as control when comparing reactions with multiple proteins.

All binding reactions were performed in 20 µL binding buffer (20 mM Tris pH 7.5, 150 mM NaCl, 1% glycerol, 0.25 mM EDTA, 2 mM MgCl_2_, 0,01% Tween-20 and 3 mM TCEP) with 10 nM labelled probe. Reactions were supplemented with 140 ng/µL fish sperm DNA (Sigma-Aldrich) for EMSAs performed with *in vitro*-produced LFY, and 200 ng/µL for EMSAs performed with recombinant 6xHis-LFY-DBD.

Binding reactions were incubated for 20 min on ice and then loaded on a 6 % native polyacrylamide gel. Gels were electrophoresed at 90 V for 75 min at 4°C and revealed with an Amersham ImageQuant 800 imager (Cytiva).

#### Recombinant protein production and purification from bacteria

6xHis-LFY-DBD was produced in *E.Coli* Rosetta2 (DE3) cells (Novagen) and purified as previously described (Hamès et al., 2008).

ASK1 was cloned into the pETM-11 expression vector (Dümmler et al., 2005), and the resulting plasmid was transformed into *E.Coli* BL21 cells (Novagen). Bacteria were grown in LB medium supplemented with kanamycin and chloramphenicol at 37 °C up to an OD_600nm_ of 0.6. Cells were then shifted to 18 °C and 0.4 mM isopropyl b-D-1-thiogalactopyranoside (IPTG) was added. After an overnight incubation, cells were sonicated in UFO buffer (25 mM Tris pH8, 150 mM NaCl, 1 mM TCEP) supplemented with one EDTA-free Pierce Protease Inhibitor Tablets (ThermoFisher). Lysed cells were then centrifuged for 30 min at 15000 rpm. Supernatant was mixed with Ni Sepharose High Performance resin (Cytiva) previously equilibrated with UFO buffer (25 mM Tris pH 8, 150 mM NaCl, 1 mM TCEP). Resin was then washed with UFO buffer containing 20 and 40 mM imidazole. Bound proteins were eluted with UFO buffer containing 300 mM imidazole and dialyzed overnight at 4 °C against UFO buffer without imidazole.

#### Recombinant protein production and purification from insect cells

The different tagged versions of ASK1, LFY and UFO were cloned in acceptor and donor plasmids (pACEBac1, pIDK and pIDS respectively; Geneva Biotech). Final acceptor plasmids containing the combination of desired coding sequences were obtained with Cre recombinase (NEB). DH10EmBacY competent cells containing the baculovirus genomic DNA (bacmid) were transformed with final acceptor plasmids. Blue-white selection was used to identify colonies with a recombinant bacmid with acceptor plasmid inserted.

Bacmid was then isolated from bacteria and mixed with X-tremeGENE HP DNA Transfection Reagent (Roche) to transfect Sf21 insect cells. 96 h after transfection, supernatant containing the recombinant baculovirus (V_0_) was collected and used to infect fresh Sf21 cells. When infected cells reached DPA (Day Post Arrest), V_1_ virus was collected. For large expression, Sf21 cells were infected with either V_1_ virus or frozen baculovirus-infected cells.

The pellet of a 0.75 L culture was sonicated in 50 mL of UFO buffer supplemented with one EDTA-free Pierce Protease Inhibitor Tablets (ThermoFisher). Sonicated cells were centrifuged for 1.5 h at 30 000 rpm, 4 °C. Supernatant was then incubated for 1 h at 4 °C with Ni Sepharose High Performance resin (Cytiva) previously equilibrated with UFO buffer. Beads were transferred into a column, and washed with 20 column volumes of UFO buffer, then UFO buffer + 50 mM imidazole. Proteins were eluted with UFO buffer containing 300 mM imidazole. Elution was dialyzed overnight at 4 °C against UFO buffer. TEV protease was added to cleave tags (0.01% w/w). When ASK1 was limiting compared to UFO, recombinant 6xHis-ASK1 from bacteria was added. The following day, elution was repassed on Dextrin Sepharose High Performance (Cytiva) and Ni Sepharose High Performance resins (Cytiva) to remove tags and contaminants.

For ASK1-UFO, ASK1-UFO-3xFLAG or UFOΔFbox-3xFLAG, proteins were concentrated with a 30 kDa Amicon Ultra Centrifugal filter (Millipore) and further purified by Size Exclusion Chromatography (SEC).

For ASK1-UFO-LFY-DBD complex purification, contaminant DNA was removed by passing proteins on Q Sepharose High Performance resin (Cytiva) pre-equilibrated with UFO buffer. Increasing salt concentrations allowed obtaining DNA-free proteins. Indicated annealed HPLC-purified oligos (see Supplemental Table2) were then added and incubated with proteins on ice for 20 min. Proteins were concentrated with a 30 kDa Amicon Ultra Centrifugal filter (Millipore) and further purified by SEC.

#### Size Exclusion Chromatography (SEC) and Size Exclusion Chromatography coupled to Multi-Angle Laser Light Scattering (SEC-MALLS)

SEC was performed with a Superdex 200 Increase 10/300 GL column (Cytiva) equilibrated with UFO buffer. Unaggregated proteins of interest were frozen in liquid nitrogen and stored at -80°C.

SEC-MALLS was performed with a Superdex 200 Increase 10/300 GL column (Cytiva) equilibrated with UFO buffer. For each run, 50 µL containing 1 mg/mL of complex was injected. Separations were performed at RT with a flow rate of 0.5 mL/min. Elutions were monitored by using a Dawn Heleos II for MALLS measurement (Wyatt Technology) and an Optilab T-rEX refractometer for refractive index measurements (Wyatt Technology). Molecular mass calculations were performed using the ASTRA software with a refractive index increment (dn/dc) of 0.185 mL/g.

#### ampDAP-seq

pTnT-5xmyc-LFY (Lai et al., 2021) was used to produce 5xmyc-LFY *in vitro* using TnT SP6 High-Yield Wheat Germ Protein Expression System (Promega). We used the ampDAP-seq libraries described in Lai et al. (Lai et al., 2021). ampDAP-seq experiments were performed in triplicates (LFY-UFO) or in duplicates (LFY_K249R_ and LFY_K249R_-UFO).

A 50 µL TnT reaction producing 5xmyc-LFY was mixed with an excess of recombinant ASK1-UFO-3xFLAG (2 µg) and 20 µL of Pierce Anti-c-Myc Magnetic Beads (ThermoScientific). DAP buffer (20 mM Tri pH 8, 150 mM NaCl, 1 mM TCEP, 0,005% NP40) was added to reach 200 µL. Mix was incubated for 1 h at 4°C on a rotating wheel. Beads were then immobilized and washed 3 times with 100 µL DAP buffer, moved to a new tube and washed once again. ampDAP-seq input libraries (50 ng) were then added, and protein-DNA mixes were incubated for 1.5 h at 4 °C on a rotating wheel. Beads were immobilized and washed 5 times with 100 µL DAP buffer, moved to a new tube and washed 2 more times. Finally, beads were mixed with 30 µL of elution buffer (10 mM Tris pH 8.5) and heated for 10 min at 90 °C.

IP-ed DNA fragments contained in the elution were amplified by PCR according to published protocol (Bartlett et al., 2017) with Illumina TruSeq primers. Remaining beads were mixed with 20 µL of 1X SDS-PAGE Protein Sample Buffer and WB were performed to check the presence of tagged proteins. PCR products were purified using AMPure XP magnetic beads (Beckman Coulter) following manufacturer’s instructions. Library molar concentrations were determined by qPCR using NEBNext Library Quant Kit for Illumina (NEB). Libraries were then pooled with equal molarity. Sequencing was done on Illumina HiSeq (Genewiz) with specification of paired-end sequencing of 150 cycles.

#### GUS staining

The different promoter versions were cloned upstream *GUS* gene in the pRB14 backbone vector (Benlloch et al., 2011). Transformants were selected with GFP seed fluorescence. The number of independent lines analyzed for each construct is indicated in each figure. GUS staining was performed on the apex of primary inflorescences of T2 plants.

Tissues were placed in ice-cold 90% acetone for 20 min at RT, and then rinsed in GUS buffer without X-Gluc (0.2% Triton X-100, 50 mM NaPO_4_ pH 7.2, 2 mM potassium ferrocyanide, 2 mM potassium ferricyanide). Tissues were transferred in GUS buffer containing 2 mM X-Gluc substrate (X-Gluc DIRECT) and placed under vacuum for 5 min. Samples were then incubated overnight at 37 °C unless specified in the legend. Finally, tissues were washed with different ethanol solutions (35%, 50%, and 70%) and pictures were taken with a Keyence VHX-5000 microscope with a VH-Z100R objective.

#### *In planta* overexpression and mutant complementation assay

Tagged versions of UFO and UFOΔFbox were cloned under the control of the *35S* promoter in pEGAD (Cutler et al., 2000). Transformants were selected with Basta treatment. Overexpressing lines with a strong gain-of-function phenotype were crossed to the strong *ufo-1* mutant. Basta-resistant F2 plants were individually genotyped to select *ufo-1 -/-* homozygous plants. For this, a fragment was amplified by PCR with oligos oGT1085 and oPR578 (see Supplemental Table 2) and digested with DpnII enzyme (NEB). Based on digestion profile, *ufo-1 -/-* plants were kept and analyzed once they reached flowering.

Mutated versions of LFY were cloned in pETH29 (Hamès et al., 2008) or pCA26 (Chahtane et al., 2013) to express LFY cDNA under the control of its endogenous promoter or the *35S* promoter, respectively. For *lfy-12* complementation assay, heterozygous *lfy-12/+* plants were transformed. Transformants were selected with GFP fluorescence and genotyped with a previously described protocol (Benlloch et al., 2011) to select *lfy-12 -/-* plants. Complementation assay was performed with T2 plants and was based on the analysis of the first 10 flowers from the primary inflorescence. Pictures were taken with a Keyence VHX-5000 microscope with a VH-Z20R objective.

#### Western Blot

For Western Blots on plant total protein extracts, indicated tissues were crushed in 2X SDS-PAGE Protein Sample Buffer (100 mM Tris pH 6.8, 20% glycerol, 2% SDS, 0.005% Bromophenol blue, and 0.8% w/v dithiothreitol) at a 1:2 w:v ratio and boiled for 5 min. Samples were then loaded on a 12% acrylamide SDS-PAGE gel. For all WB, transfer was performed with iBlot2 Dry Blotting System (Invitrogen) using default parameters. Membranes were blocked for 1 h at RT with 5% milk TBST and then incubated overnight at 4 °C with 5% milk TBST solution containing antibody (1:1000 for anti-FLAG and 1:5000 for anti-myc). Revelation was performed with Clarity Western ECL substrate (Bio-Rad). Pictures were taken with a ChemiDoc MP Imaging System (BioRad).

#### Cryo-EM sample preparation, data collection and data processing

An aliquot of the SEC-purified ASK1-UFO-LFY-LUBS1 complex was thawed on ice (see Supplemental Table 2 for DNA sequence). Subsequently, 3.5 μl of the complex at 1 mg/mL were deposited onto glow-discharged (25 mA, 30 s) C-flat Au grid R 1.2/1.3 300 mesh (Electron Microscopy Sciences), blotted for 5.5 s with force 0, at 20°C and 100% humidity using a Mark IV Vitrobot (FEI, Thermo Fisher Scientific) and plunge-frozen in liquid ethane for specimen vitrification. A dataset of about 1’000 movies of 40 frames was acquired on a 200 kV Glacios (Thermo Fisher Scientific) electron microscope (Supplemental Table 3) at a nominal magnification of 36’000 with a physical pixel size of 1.145 Å.

The raw movies, acquired with SerialEM on a Gatan K2 Summit camera (Supplemental Table 3), were imported to Cryosparc live (Punjani et al., 2017) for motion correction and CTF estimation. The dose-weighted micrographs were used for particle picking with crYOLO 1.7.6 and the general model for low-pass filtered images (Wagner et al., 2019). Particle coordinates were imported to Cryosparc, where all subsequent steps were performed. After manual inspection, a subset of 761 micrographs was selected based on CTF fit resolution, total and per frame motion, average defocus and relative ice thickness. A raw particle stack of 282’567 images was extracted at 256x256 pixels² box size, binned twice and subjected into 2D classification to remove false positive picks. 207’392 particles from the selected class averages were re-extracted, re-centered at full size and submitted for a second round of 2D classification. All class averages showing clear protein features were selected and the resulting 147’849 particles were used for *ab initio* reconstruction with 3 classes and subsequent heterogeneous refinement of the resulting volumes. Of those 3 classes, 2 looked like a protein-DNA complex with the most apparent difference being the presence or not of an extra electron density at one edge of the DNA helix. The last class had no recognizable features and was used as a decoy to remove “junk” particles. Each subset and volume of the 2 first classes was refined separately with Non-Uniform refinement (Punjani et al., 2020) resulting into 2 distinct reconstructions of about 4.2 Å resolution, where the DNA model, the crystal structure of LFY-DBD and the AlphaFold2 models of UFO and ASK1 could be unambiguously fitted into the electron density. The second of these classes could fit a LFY-DBD dimer, while in the first class there was density only for the LFY-DBD molecule that directly interacts with UFO (Figure S6). The unsharpened maps of each reconstruction were used for post-processing with DeepEMhancer (Sanchez-Garcia et al., 2021). Figures were prepared with Chimera (Pettersen et al., 2004) or ChimeraX (Pettersen et al., 2021).

#### Cryo-EM model building

Ideal B-form DNA was generated in Coot (Emsley et al., 2010) and then manually built into the electron density. The resulting model was further refined using phenix.real_space_refine (Afonine et al., 2018). A single monomer of LFY-DBD was manually placed in the electron density, followed by fitting in ChimeraX (Pettersen et al., 2021). The biological LFY-DBD dimer was then downloaded from the RCSB PDB (Hamès et al., 2008; entry 2VY1) and used as a guide to place the second LFY monomer, followed by fitting to density in ChimeraX. Alphafold models (Jumper et al., 2021) of ASK1 (uniprot ID: Q39255) and UFO (uniprot ID: Q39090) were both downloaded from the EBI, preprocessed to remove low confidence regions in phenix.process_predicted_model (Terwilliger et al., 2022), then placed manually and then fit to density in ChimeraX.

#### Bioinformatic analyses

##### Read mapping and peak calling

Reads processing and peak calling of LFY, LFY-UFO, LFY_K249R_ and LFY_K249R_-UFO ampDAP-seq data were performed as previously published (Lai et al., 2020). Briefly, the quality of sequencing data was analyzed with fastQC v0.11.7 and adapters were removed with NGmerge v0.2_dev (Gaspar, 2018). Bowtie2 v2.3.4.1 was used for mapping to the TAIR10 *A. thaliana* reference genome (Berardini et al., 2015). Reads mapped to a single location and with maximum two mismatches were retained Duplicates were removed with the samtools dedup program v1.8.

Bound regions (i.e. peaks) were identified with MACS2 v2.2.7.1, using input DNA from Lai et al. as control (Lai et al., 2021). Consensus peaks were selected with MSPC v4.0.0 (Jalili et al., 2015) by retaining peaks called in all replicates, and resizing them by ±200 bp around the peak maximum for further analysis.

##### Analyses of ampDAP-seq experiments

To compare binding in different experiments, peaks were merged according to a previously published procedure (Lai et al., 2020). Bound peaks were considered as common if they overlapped by at least 80%, while the remaining non-overlapping portion of either peak was < 50%. Peaks that did not overlap by at least 50% were considered as new peaks. The same procedure was used to assess experimental reproducibility (comparisons between replicates of the same experiment), where peaks were normalized by the number of reads mapped in library (RPKM).

As the fraction of reads mapped in peaks is much lower for LFY than LFY-UFO ampDAP-seq (∼25% vs ∼40%, respectively), normalizing reads count by all reads mapped along the genome would introduce a bias and estimate the LFY relative coverage (RPKM) towards lower values compared to LFY-UFO. In addition to this consideration, experimental proof from EMSAs suggests that UFO does not strongly affect binding intensity of the complex at canonical LFYBS (which represent most peaks). Hence, reads count at each peak was normalized by the total number of reads mapped within all LFY and LFY-UFO merged peaks. Then, the mean normalized coverage from each experiment, divided by the peak size, was computed for each peak. The same strategy was applied when comparing LFY_K249R_ and LFY_K249R_-UFO (Figure 5B), LFY_K249R_ and LFY (Figure S5H) and LFY, LFY-UFO, LFY_K249R_ and LFY_K249R_-UFO (Figure 5C).

The Coverage Fold Change (CFC) was computed on merged peaks as the ratio between mean normalized peak coverage in LFY-UFO and LFY (Figure 3B) or mean normalized coverage in LFY_K249R_-UFO and LFY_K249R_ (Figure 5B).

##### Motif search in bound regions

Merged peaks of LFY and LFY-UFO datasets were sorted based on decreasing CFC value. The top 600 peaks (i.e. highest CFC values) were used for a motif search using MEME-ChIP v4.12.0 using options - nmeme 600 -meme-maxsize 600*1000 -meme-nmotifs 1 -dreme-m 0 -noecho and the JASPAR 2018 core plants non-redundant database (Machanik and Bailey, 2011). For dLUBS, we used options -meme-minw 20 -meme-maxw 30, while for mLUBS we used -meme-minw 16 -meme-maxw 19. To retrieve the LFY motif in Figure 3C the 600 LFY ampDAP-seq peaks with strongest coverage were fed to MEME-ChIP with options -nmeme 600 -meme-nmotifs 1 --meme-minw 19 -meme-maxw 19 –pal.

##### Receiver Operating Characteristics analysis

From the dataset of merged peak set (peaks found in LFY or in LFY-UFO experiments or in both), peaks were sorted based on decreased CFC value, the top 20% peaks were selected, and among these, the first 600 used for motif determination were excluded to avoid overfitting, for a total of 3243 final peaks. A negative set of the same size was created using a previously published method, which allows searching for sequences from the *A. thaliana* genome (TAIR10 reference) with the same GC content and genomic origin as the positive set (Stigliani et al., 2019). Both sets were scanned with dLUBS and mLUBS PWMs as well as with the LFY PWM with dependencies as published in (Moyroud et al., 2011) using an in-house script available on our GitHub page. The ROC plot was then created with the R ‘plotROC’ package v2.2.1.

##### LFY in dLUBS within LFY-UFO-specific regions vs LFY in LFY-specific regions

To assess whether the scores of LFYBS within dLUBS were comparable to the scores of canonical LFYBS, we used the peaks from the comparison of LFY vs LFY-UFO ampDAP-seq and resized them (+/-50 bp around the peak maximum). We used the dLUBS matrix to scan the resized sequences and retained the best site per sequence. We then retrieved sequences corresponding to the dLUBS site and computed the score of the LFYBS present in dLUBS using the LFY PWM (Moyroud et al., 2011). The values obtained in the 20% most LFY-UFO-specific sequences (20% highest CFC) is shown in the boxplot. The 20% lowest CFC peaks were scanned with the LFY PWM (Moyroud et al., 2011) to generate the box-plot in Figure S3D.

##### Microarray data analysis

Microarray data were retrieved from AtGenExpress (Schmid et al., 2005) for inflorescence tissue in the *ufo* (ATGE_52A-C) vs Col-0 background (ATGE_29A-C). The ‘gcrma’ R package was used to adjust probe intensities and convert them to expression measures, and then the ‘limma’ package was used to fit the model and smooth standard errors. A Benjiamini-Hochberg correction was applied to p-values and fold change (FC) was computed as the ratio between expression in wt versus the *ufo* mutant. Only genes with |log2(FC)|> 0.5 and adjusted p-value < 0.05 were considered as significantly differentially expressed.

##### ChIP-seq datasets and analysis of ChIP-seq vs ampDAP-seq

We collected the raw data of all available LFY ChIP-seq datasets: GSE141704 (Jin et al., 2021), GSE96806 (Goslin et al., 2017), GSE64245 (Sayou et al., 2016), GSE24568 (Moyroud et al., 2011). Mapping and peakcalling analysis were performed with the same procedure as ampDAP-seq, except that peaks were resized to 600 bp around the peak maximum, and the –q option of MACS2 was set to 0.1. Coverage of the resulting peaks was calculated as the average of normalized read coverage for each replicate. Peaks from the four datasets were merged through a four-way comparison following the same procedure used for ampDAP-seq. Bedtools intersect (v2.30.0) was used with options -wa –f 0.8 -F 0.8 -e to find the peaks common to the merged ChIP-seq peaks and the 20% most LFY-UFO-specific genomic regions (highest CFC value from ampDAP-seq). Peaks were assigned to genes by extending gene regions 3 kb upstream of the TSS and 1 kb downstream of the TTS and using bedtools intersect (options -f 0.8 -F 0.8 –e) to identify genes in the vicinity of peaks. The bound genes obtained were crossed with the list of differentially expressed genes in *ufo* inflorescences.

##### Identification of the URM from published LFY ChIP-seq data

To test whether the URM could be identified *de novo* (Figure S3E), we collected the 298 regions bound by LFY ChIP-seq data of inflorescence tissue (Goslin et al., 2017) for which the binding intensity was twice greater *in vivo* relative to *in vitro* (LFY ampDAP-seq). We resized these regions +/- 55 bp around the ChIP-seq peak maximum. The corresponding sequences were searched with the LFY PWM (Moyroud et al., 2011) to identify all LFYBS with a PWM score > -23. Assuming that a recruiting motif should be at a fixed distance from the LFYBS, we created 140 batches, corresponding to sequences with size ranging from 4 to 10 bp, distant from 1 to 20 bp at both sides of the canonical LFYBS. Each of the 140 batches of sequences was used as input with MEME-ChIP for motif discovery with the motif size constrained to the length of the sequences in a given batch.

#### QUANTIFICATION AND STATISTICAL ANALYSES

All DLRA data were analyzed using R Studio software and are presented as mean ± SD. All statistical methods are indicated within the figure legends. One-way ANOVA was used to analyze experimental data with more than two experimental groups. Welch’s ANOVA was performed when the homogeneity of variance assumption was not met. Two-tailed unpaired Student’s t-test was used for other data analyses.

## SUPPLEMENTAL ITEM TITLES

**Supplemental Table 1: List of plasmids used in this study**

**Supplemental Table 2: List of oligonucleotides used in this study**

**Supplemental Table 3: Data collection and refinement statistics**

**Supplemental Item 1: Uncropped gels (Western Blots and EMSAs)**

---

**Figure S1.**
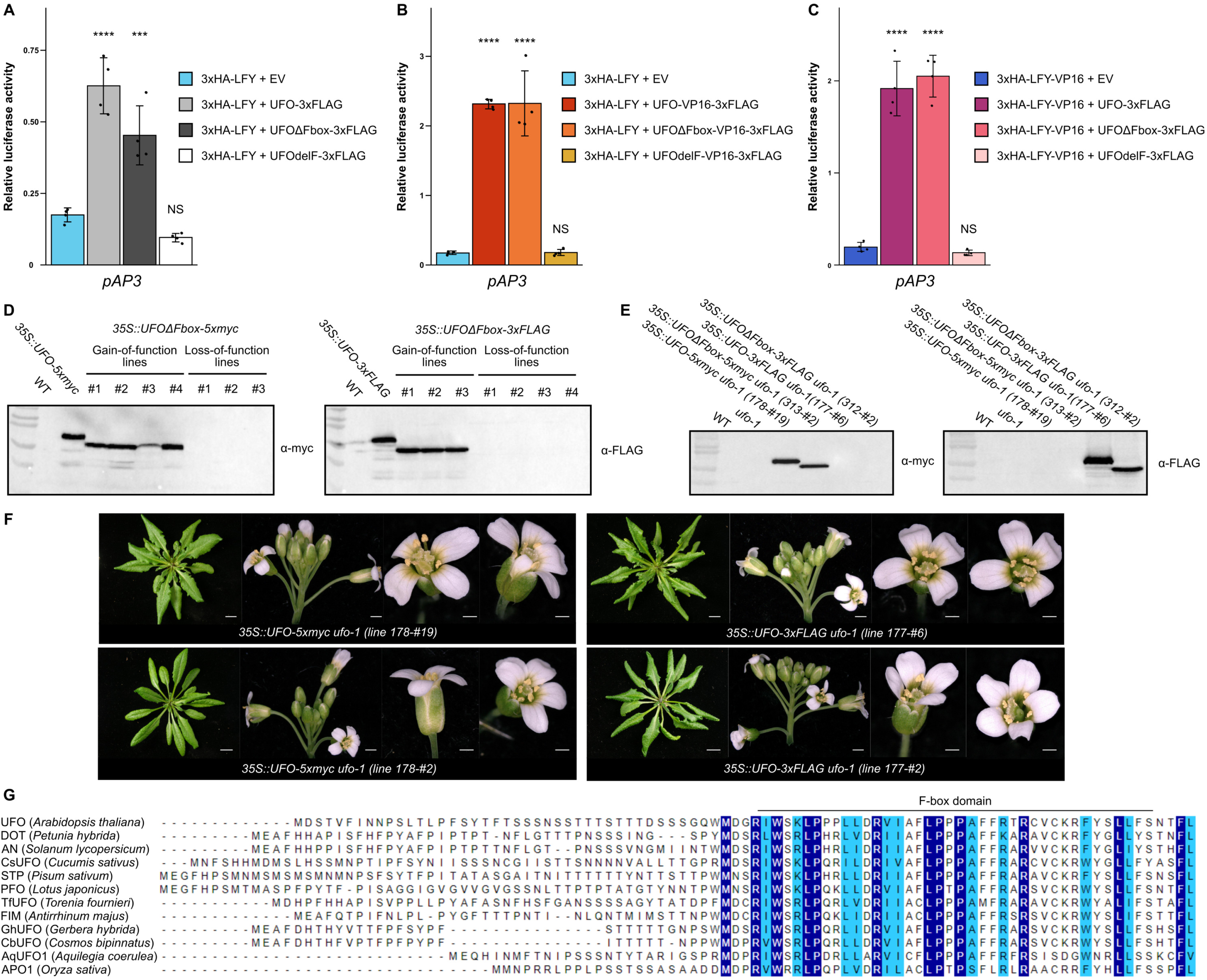
UFO has SCF-dependent and independent functions, related to Figure 1. (A to C) *pAP3* activation measured by DLRA in Arabidopsis protoplasts. EV = Empty Vector (pRT104-3xHA). UFOΔFbox corresponds to a deletion of the whole N-terminal part comprising the F-box domain (aa. 1-90), while UFOdelF corresponds to an internal deletion in the F-box domain (aa. 50-62; Risseeuw et al., 2013). Data represent averages of independent biological replicates and are presented as mean ± SD, each dot representing one biological replicate (n = 4). One-way ANOVA with Tukey’s multiple comparisons test. Stars above bars represent a significant statistical difference compared to 3xHA-LFY + EV or 3xHA-LFY-VP16 + EV negative controls (NS: p > 0.05,*: p < 0.05, **: p < 0.01, ***: p < 0.001 and ****: p < 0.0001). (D) Western Blot on protein extracts from T1 plants from different phenotypic classes described in Figure 1F. *35S::UFO-5xmyc* (178-#19) and *35S::UFO-3xFLAG* (177-#6) plants were used as positive controls. Total proteins were extracted from rosette leaves. Note the difference of molecular weight between UFO and UFOΔFbox. Loss-of-function defects are likely due to silencing of both transgene-encoded *UFOΔFbox* and endogenous *UFO.* Gels were cropped (see Supplemental Item 1). (E) Western Blot on protein extracts from F2 plants described in Figure 1H and S1F. Total proteins were extracted from rosette leaves. Gels were cropped (see Supplemental Item 1). (F) *ufo-1* complementation assay with *35S::UFO* lines. Rosette leaves (right, scale bar, 1 cm), inflorescence (middle, scale bar 1 mm) and flower (right, scale bar, 0.5 mm) phenotypes are shown. Primary inflorescences were removed to observe rosette phenotype. For each construct, two lines were crossed to *ufo-1* and at least 5 plants were analyzed per line. As in Risseeuw et al, our *35S::UFO* lines displayed relatively milder phenotypes than the *35S::UFO* phenotypes reported by Lee et al. (Lee et al., 1997; Risseeuw et al., 2013). Note that the *35S::UFO-5xmyc* 178-#2 line did not display the serrated leaves phenotype. (G) Sequence alignment of UFO N-terminal region. The F-box domain is represented (Gagne et al., 2002). In selected species, presented proteins were identified as *UFO* homologs and their role was confirmed genetically (Chen et al., 2021; Ikeda et al., 2005; Levin and Meyerowitz, 1995; Li et al., 2019; Lippman et al., 2008; Sasaki et al., 2012; Sharma et al., 2019; Simon et al., 1994; Souer et al., 2008; Taylor et al., 2001; Zhang et al., 2003; Zhao et al., 2016).

**Figure S2.**
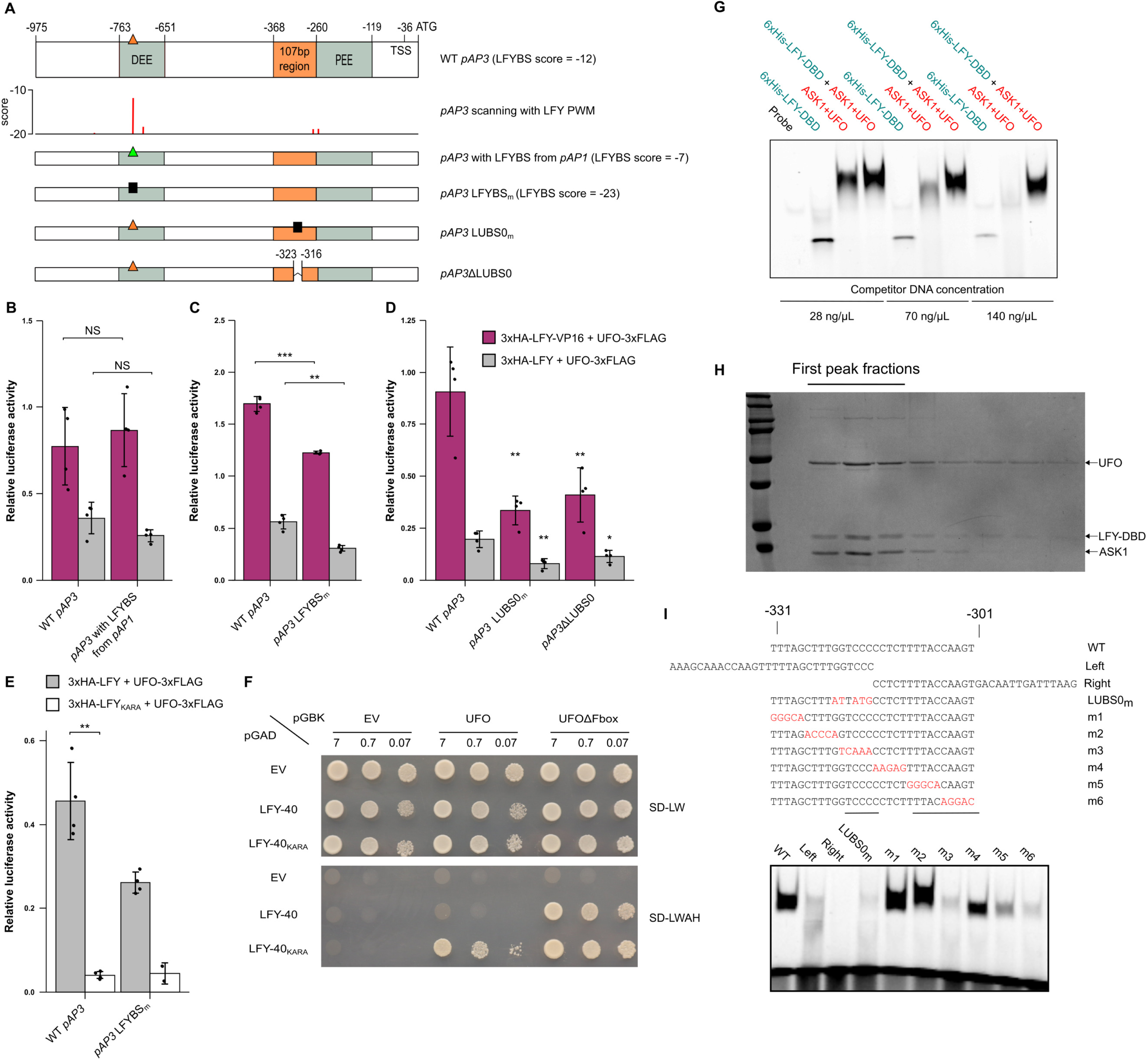
Analysis of *pAP3* activation by LFY-UFO, related to Figure 2. (A) Schematic representation of *pAP3*. Top row represents WT *pAP3,* the second row represents the scores for the best LFYBS obtained by scanning WT *pAP3* sequence with LFY PWM (the best binding sites correspond to the less negative score values; Moyroud et al., 2011). Other rows represent the different *pAP3* versions used in (B-E). LFYBS mutation corresponds to the previously described *site1m-site2m* mutation (Lamb et al., 2002). The LUBS0 mutation is described in (H). (B-E) *pAP3* activation with promoter versions described in (A) and indicated effectors. Note that in (E) only n = 2 biological replicates were done for 3xHA-LFY_KARA_ + UFO-3xFLAG on *pAP3* LFYBS_m_ (no statistical test). (F) Effect of the LFY_KARA_ mutation on LFY-UFO interaction in Yeast-Two-Hybrid (Y2H). EV = Empty Vector. LFY-40 is a LFY version lacking the first 40 aa and better tolerated by yeast cells. Values correspond to the different dilutions (OD = 7, 0.7 and 0.07). Top picture corresponds to the non-selective plate lacking Leucine and Tryptophan (SD -L-W), and bottom picture to the selective plate lacking Leucine, Tryptophan, Histidine and Adenine (SD -L-W-A-H). Pictures were taken at day + 4. (G) EMSA with ASK1-UFO, LFY-DBD and LUBS0 DNA probe. Different competitor DNA concentrations were tested as indicated. Gel was cropped and only protein-DNA complexes are shown (see Supplemental Item 1). (H) Coomassie-stained SDS-PAGE gel of the different SEC-MALLS fractions from Figure 2F. Each lane corresponds to a 0.5 mL fraction. (I) EMSA with ASK1-UFO, LFY-DBD and indicated DNA probes. Sequences with coordinates relative to *AP3* start codon (top). Red letters indicate mutated bases. Bars under sequences represent the regions required for ASK1-UFO-LFY-DBD binding. EMSA with described DNA probes (bottom). Each DNA probe was mixed with the same ASK1-UFO-LFY-DBD protein mix. Note that the LUBS0 mutation also reduced *pAP3* activation in protoplasts (Figure S2D). For bar charts, data represent averages of independent biological replicates and are presented as mean ± SD, each dot representing one biological replicate (n = 4 unless specified). One-way ANOVA with Tukey’s multiple comparisons tests (D). In (D), one-way ANOVA was performed with data from the same effector, and stars above bars represent a significant statistical difference compared to WT *pAP3*. Unpaired t-tests (B, C and E). (NS: p > 0.05,*: p < 0.05, **: p < 0.01, ***: p < 0.001 and ****: p < 0.0001).

**Figure S3.**
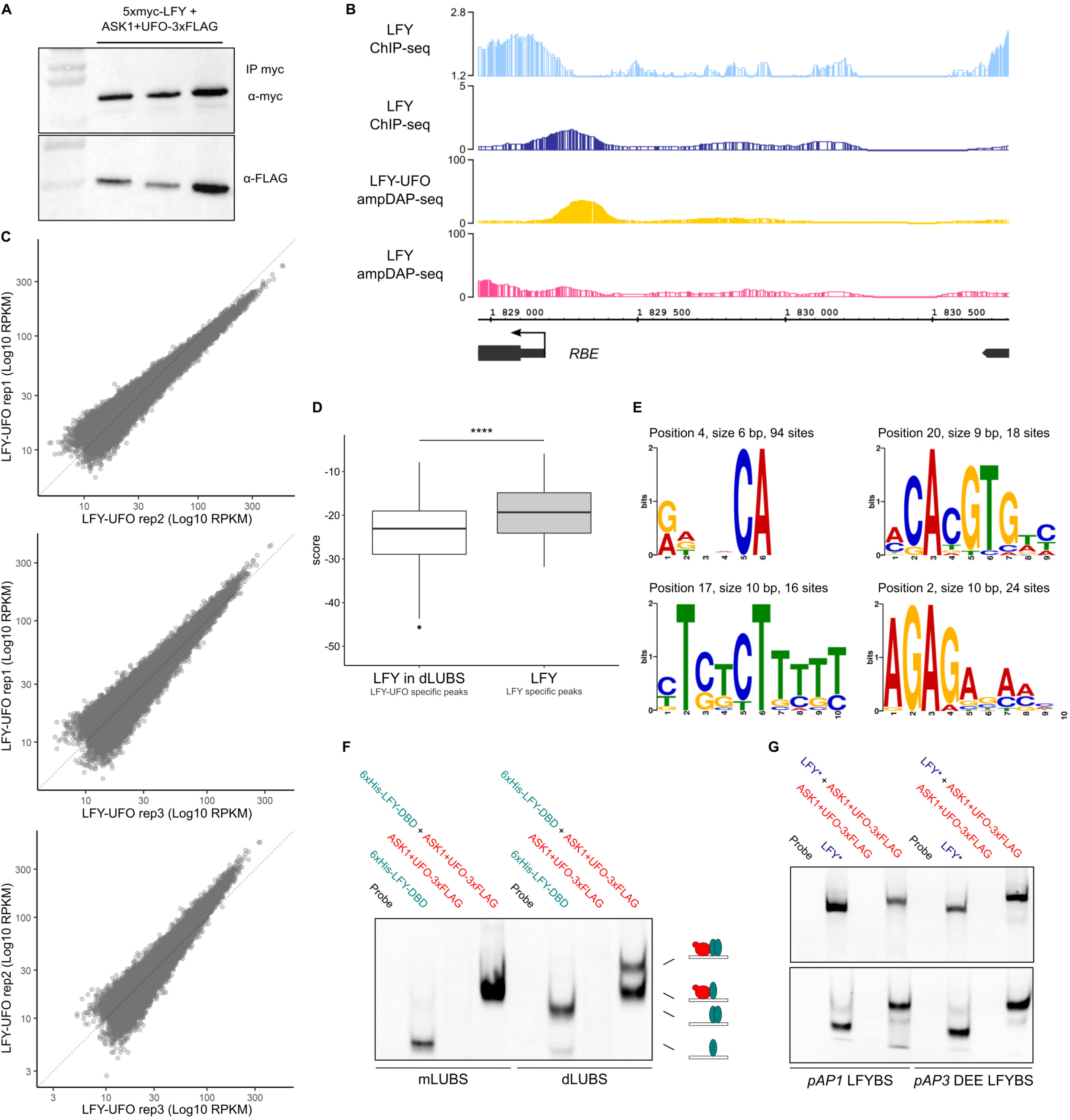
Genome-wide analysis of LFY-UFO DNA binding, related to Figure 3. (A) Western Blot after DNA elution during ampDAP-seq experiment. After DNA elution, 20 µL of 1X SDS-PAGE Protein Sample Buffer was added to the remaining beads to run WB. Each lane represents one replicate. Gels were cropped (see Supplemental Item 1). (B) IGB view of *pRBE* showing LFY ChIP-seq in inflorescences (light blue; Goslin et al., 2017) or seedlings (dark blue; Sayou et al., 2016), LFY-UFO ampDAP-seq (yellow; this study), LFY ampDAP-seq (pink; Lai et al., 2021), numbers indicate read number range. (C) Assessment of experimental reproducibility of ampDAP-seq experiment through the comparison of replicates datasets 2 by 2. (D) Score distribution of LFY PWM with dependencies (Moyroud et al., 2011) within dLUBS (best site on 20% most LFY-UFO-specific genomic regions, high CFC) and in canonical LFYBS (best site on 20% most LFY-specific genomic regions, low CFC). Best sites were selected within ±25 bp around the peak maximum (see Methods). Wilcoxon rank sum test (****: p < 0.0001). (E) *De novo* identification of URM from LFY ChIP-seq data. The panel shows the motifs identified at a fixed distance from LFY canonical binding sites in 298 regions harboring high LFY ChIP-seq to LFY ampDAP-seq coverage ratio. The text above each motif gives the motif’s start position relative to the canonical LFYBS, its length and the number of sites used to build the motif. (F) EMSA with mLUBS and dLUBS highest score sequences. 6xHis-LFY-DBD and ASK1-UFO-3xFLAG complex are recombinant. Drawings represent the different types of complexes involving LFY-DBD (pale blue) and ASK1-UFO (red) on DNA. LFY-DBD binds as a monomer as previously reported (Hamès et al., 2008). Gel was cropped and only protein-DNA complexes are shown (see Supplemental Item 1). (G) EMSA with DNA probes corresponding to *pAP1* and *pAP3* DEE LFYBS. LFY* refers either to *in vitro*-produced 5xmyc-LFY (top) or recombinant 6xHis-LFY-DBD (bottom). Note that probes used here have the same length as those used to study LUBS. Gels were cropped and only protein-DNA complexes are shown. Gels were cropped and only protein-DNA complexes are shown (see Supplemental item 1).

**Figure S4.**
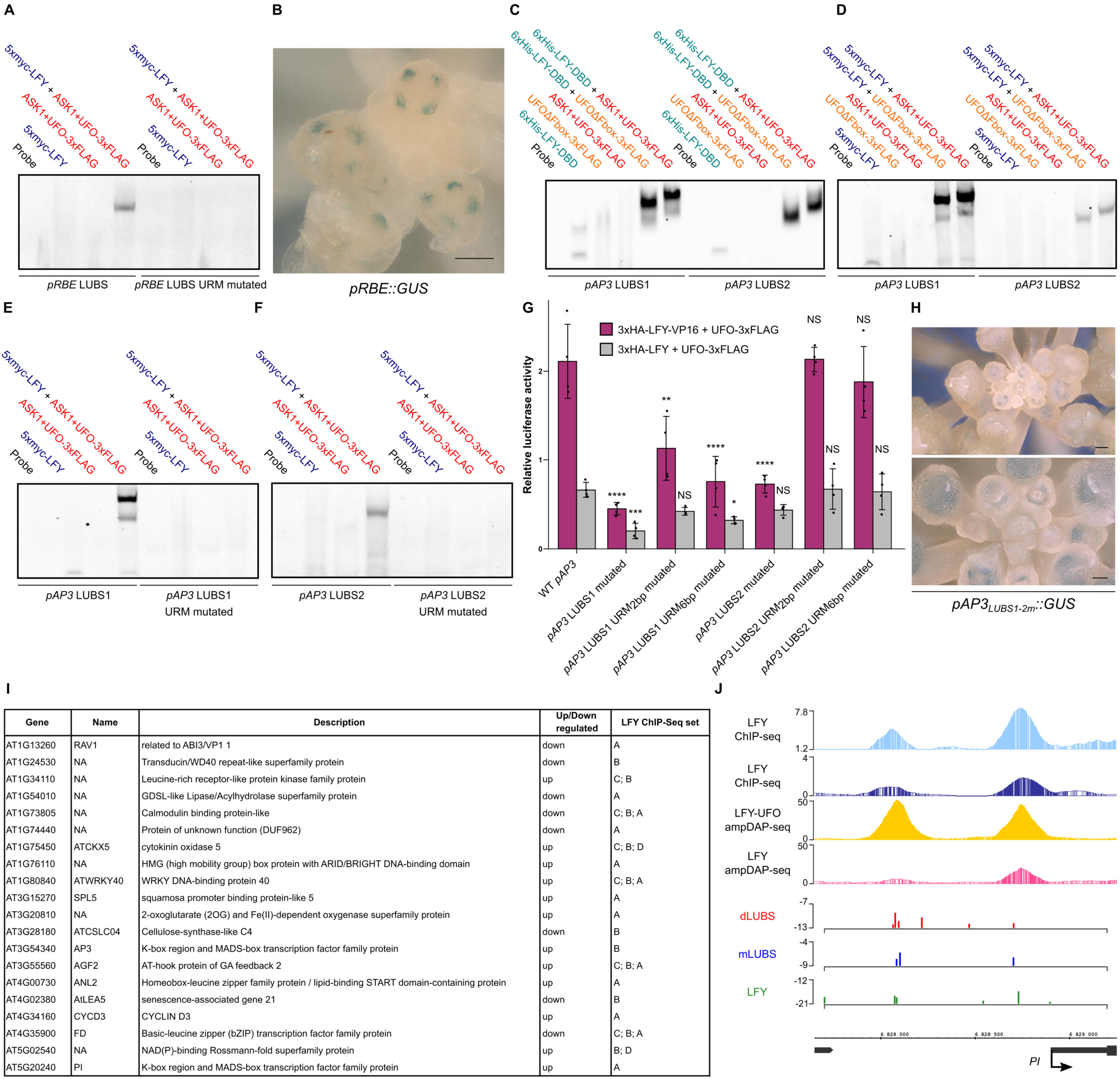
Characterization of *pRBE* and *pAP3* LUBS, related to Figure 4. (A) EMSA with probes corresponding to *pRBE* LUBS, WT or with URM mutated. (B) *In vivo* analysis of *pRBE::GUS* fusions. Same as in Figure 4C, with another view showing staining in the four petal primordia (scale bar, 50 µm). (C-D) EMSA with *pAP3* LUBS1 and LUBS2 DNA probes and recombinant 6xHis-LFY-DBD (C) or *in vitro*-produced 5xmyc-LFY (D). Note the difference of complex size between UFO and UFOΔFbox. (E-F) EMSA with DNA probes corresponding to *pAP3* LUBS1 (E) and LUBS2 (F), WT or with URM mutated. (G) Promoter activation measured by DLRA in Arabidopsis protoplasts with indicated effectors. Different promoter versions were tested as indicated under x-axis. Either 2 bp (high-informative CA) or 6 bp (whole URM) of *pAP3* LUBS1 and LUBS2 URM were mutated. Data represent averages of independent biological replicates and are presented as mean ± SD, each dot representing one biological replicate (n = 4). One-way ANOVA with Tukey’s multiple comparisons tests. One-way ANOVA were performed with data from the same effector and stars represent a statistical difference compared to WT *pAP3* promoter. (NS: p > 0.05,*: p < 0.05, **: p < 0.01, ***: p < 0.001 and ****: p < 0.0001). (H) *In vivo* analysis of *pAP3_LUBS1-2m_::GUS* fusions. Same as in Figure 4F, except that staining incubation time was increased to 17 h. Representative pictures are shown (top scale bar, 100 µm, bottom scale bar, 50 µm). The faint *AP3* pattern suggests that other LUBS (such as LUBS0) may take over but less efficiently. Note that with this staining incubation time, all plants expressing *pAP3::GUS* showed a highly saturated staining. (I) List of candidate LFY-UFO target genes selected as i) present in regions specifically bound by LFY-UFO in ampDAP-seq (high CFC) ii) bound *in vivo* in LFY ChIP-seq experiments (Goslin et al., 2017 (A); Sayou et al., 2016 (B); Moyroud et al., 2011 (C); Jin et al., 2021 (D)) and iii) deregulated in *ufo* inflorescences (Schmid et al., 2005). (J) From the top: IGB view of *PISTILLATA* promoter region showing LFY ChIP-seq in inflorescences (light blue; Goslin et al., 2017) or seedlings (dark blue; Sayou et al., 2016), LFY-UFO ampDAP-seq (yellow; this study), LFY ampDAP-seq (pink; Lai et al., 2021), numbers indicate read number range. Below, predicted binding sites using the dLUBS, mLUBS models from Figure 3C and LFY PWM with dependencies (Moyroud et al., 2011), y-axis represents score values. For all EMSAs, gel pictures were cropped and only protein-DNA complexes are shown (see Supplemental Item 1).

**Figure S5.**
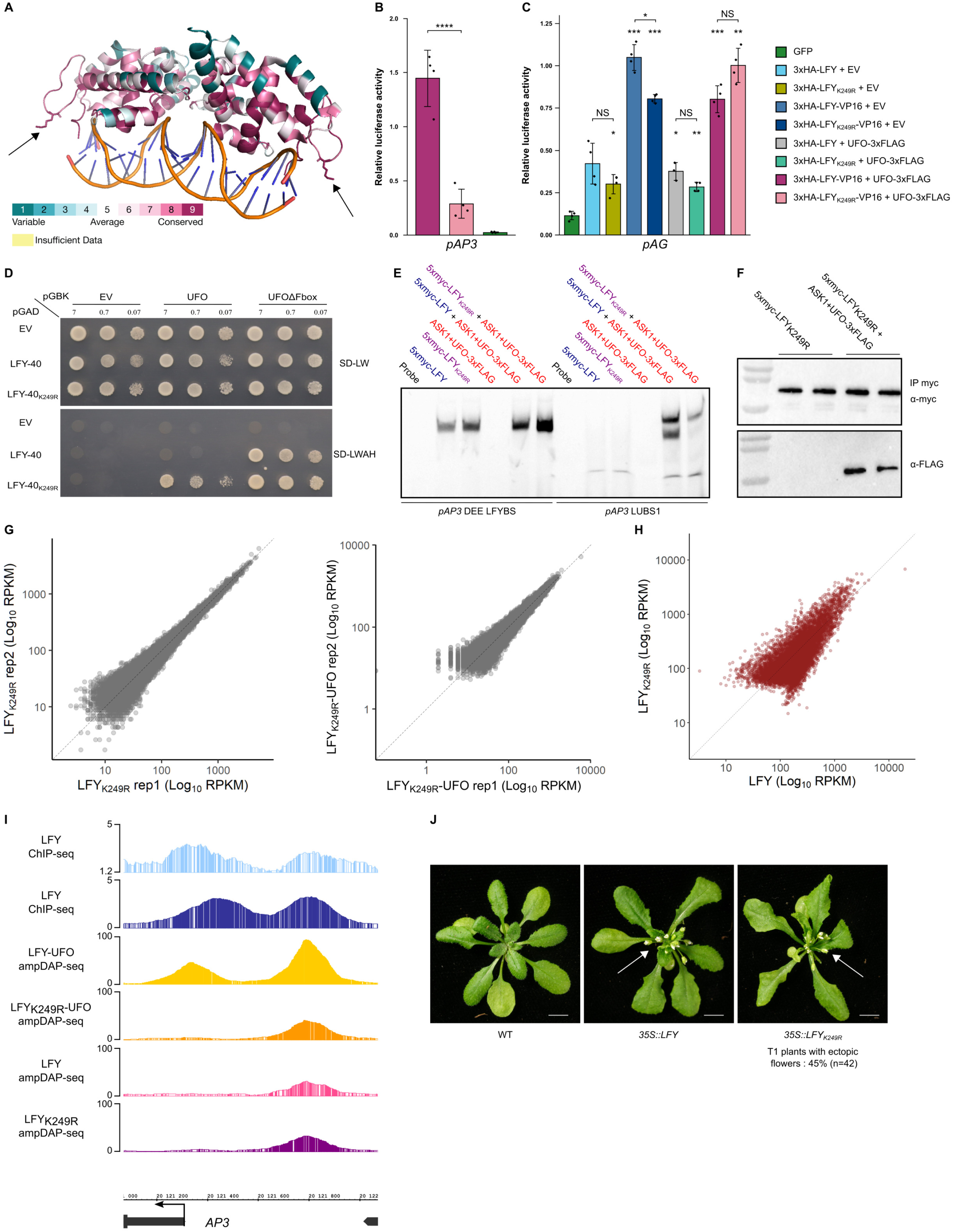
The LFY K249 is essential for LFY-UFO-LUBS complex formation, related to Figure 5. (A) Structure of LFY-DBD (Hamès et al., 2008). Residues were colored by conservation using Consurf with default parameters (Ashkenazy et al., 2016). K249 residues on each LFY monomer are represented as sticks and indicated with arrows. Note that the K249-containing loop is highly conserved. (B-C) Promoter activation measured by DLRA in Arabidopsis protoplasts with indicated effectors (right). EV = Empty Vector (pRT104-3xHA). Tested promoters are indicated below each graph. Note that for 3xHA-LFY+UFO-3xFLAG on *pAG* only n = 3 biological replicates are shown. Data represent averages of independent biological replicates and are presented as mean ± SD, each dot representing one biological replicate (n = 4 unless specified). One-way ANOVA with Tukey’s multiple comparisons tests (B) or Welch’s ANOVA with Games-Howell post-hoc test (C). In (C), stars above bars represent a statistical difference compared to GFP. (NS: p > 0.05,*: p < 0.05, **: p < 0.01, ***: p < 0.001 and ****: p < 0.0001) (D) Effect of the LFY_K249R_ mutation on LFY-UFO interaction in Y2H. EV = Empty Vector. LFY-40 is a LFY version lacking the first 40 aa and better tolerated by yeast cells. Values correspond to the different dilutions (OD = 7, 0.7 and 0.07). Top picture corresponds to the non-selective plate lacking Leucine and Tryptophan (SD -L-W), and bottom picture corresponds to the selective plate lacking Leucine, Tryptophan, Histidine and Adenine (SD -L-W-A-H). Pictures were taken at day + 4. (E) EMSA with DNA probes corresponding to *pAP3* DEE LFYBS and *pAP3* LUBS1 and indicated proteins. *pAP3* DEE LFYBS DNA probe was used as a control for binding on canonical LFYBS. Gel was cropped and only protein-DNA complexes are shown (see Supplemental Item 1). (F) WB after DNA elution during ampDAP-seq experiment. After DNA elution, 20 µL of 1X SDS-PAGE Protein Sample Buffer was added to the remaining beads to run WB. Each lane represents one replicate. Gels were cropped (see Supplemental Item 1). (G) Reproducibility of ampDAP-seq experiments with LFY_K249R_ (left) and LFY_K249R_-UFO (right) through the comparison of replicates datasets 2 by 2. (H) Comparison of peak coverage in LFY_K249R_ (y-axis, this study) and LFY (x-axis, Lai et al., 2021) ampDAP-seq experiments. (I) Integrated Genome Browser (IGB) view of *pAP3* showing LFY ChIP-seq in inflorescences (light blue; Goslin et al., 2017) or seedlings (dark blue; Sayou et al., 2016), LFY-UFO ampDAP-seq (yellow; this study), LFY_K249R_-UFO ampDAP-seq (orange; this study), LFY ampDAP-seq (pink; Lai et al., 2021), and LFY_K249R_ ampDAP-seq (purple; this study). Numbers indicate read number range. (J) Pictures of WT and representative transgenic plants expressing *35S::LFY* or *35S::LFY_K249R_* (scale bar, 1 cm). The white arrows indicate ectopic rosette flowers. *35S::LFY* was obtained previously (Sayou et al., 2016). 42 T1 plants expressing *35S::LFY_K249R_* were analyzed; the percentage of plants with a LFY overexpressing phenotype is comparable to the one obtained with *35S::LFY* (Sayou et al., 2016).

**Figure S6.**
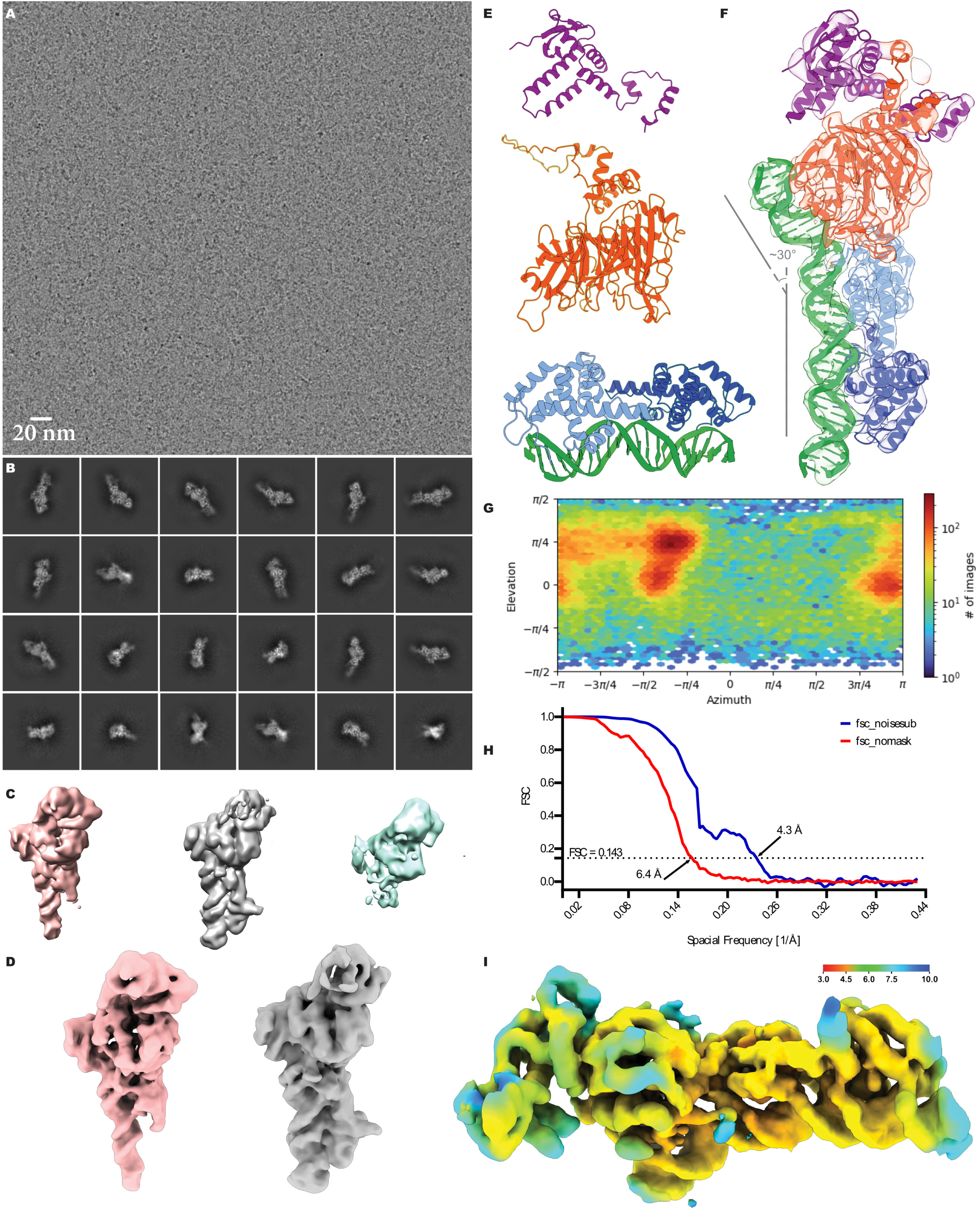
UFO binds DNA and LFY, related to Figure 6. (A) A representative micrograph of the ASK1-UFO-LFY-DNA complex in vitreous ice. (B) Selected 2D class averages of the particles submitted to *ab initio* reconstruction and heterogeneous refinement for 3D classification. (C) Intermediate reconstructions of the 3D classes after heterogeneous refinement. (D) Final reconstructions of ASK1-UFO-LFY-DNA complexes (involving either a LFY-DBD monomer (pink) or a LFY-DBD dimer (gray)) after Non-Uniform refinement. (E) Unprocessed AlphaFold2 model for ASK1 (top, purple; uniprot ID, Q39255), UFO (middle, red; uniprot ID, Q39090) and the LFY-DBD dimer/DNA crystallographic structure (bottom, pale and dark blue for the LFY-DBD dimer and green for the DNA; PDB entry: 2VY1). (F) Cryo-EM density map color-coded by fitted molecule. Note the kink on DNA induced by the presence of UFO. (G) Heat map of the angular distribution of particle projections contributing for the final reconstruction of the complete ASK1-UFO-LFY-DNA complex (with a LFY-DBD dimer). (H) Gold-standard Fourier shell correlation (FSC) curves. The dotted line represents the 0.143 FSC threshold, which indicates a nominal resolution of 6.4 Å for the unmasked (red) and 4.3 Å for the masked (blue) reconstruction. (I) View of the post-processed map of the complete ASK1-UFO-LFY-DNA complex, colored according to the local resolution.

