## Supplemental figures for "The F-box cofactor UFO redirects the LEAFY floral regulator to novel *cis*-elements"

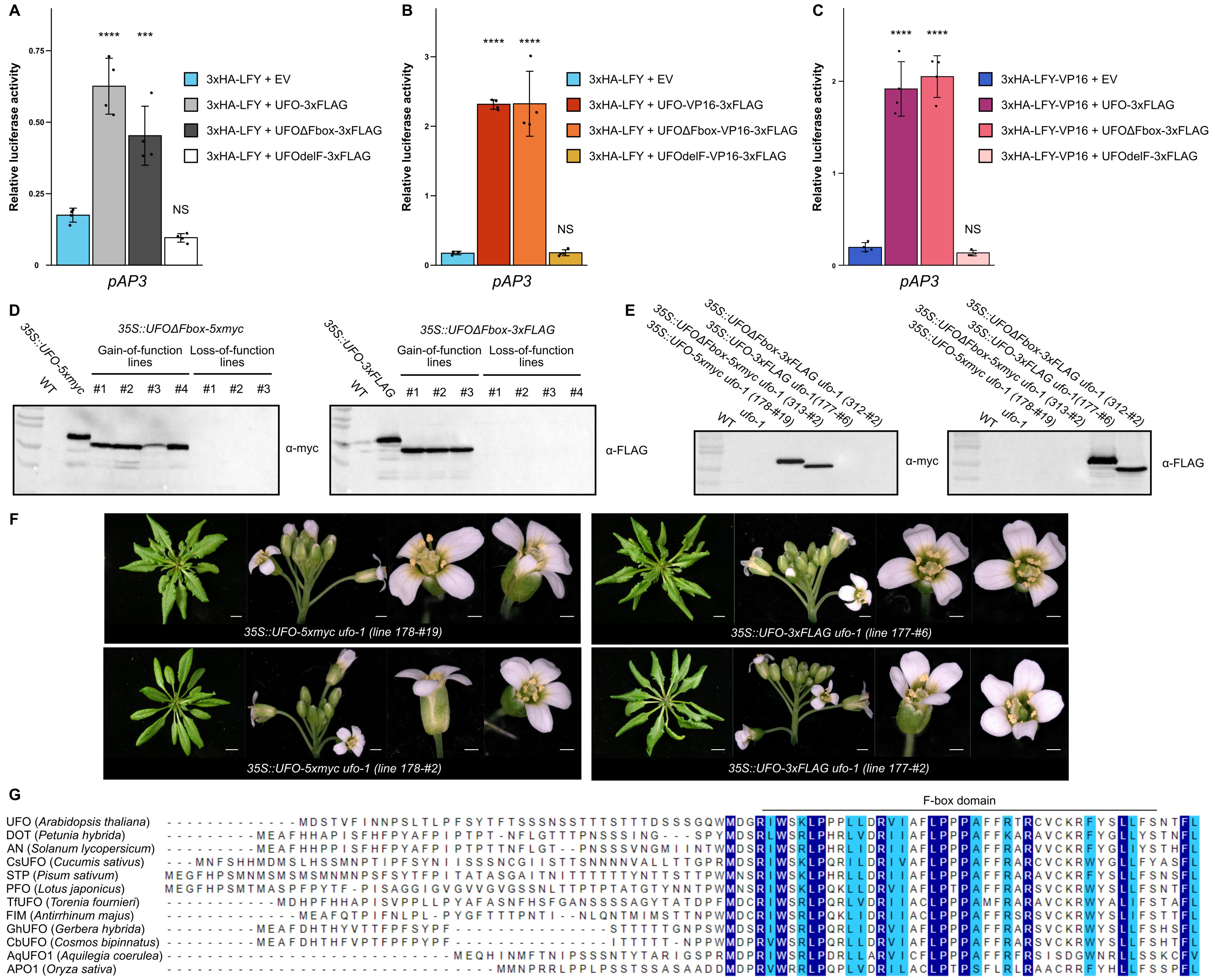
A
B
C
Relative luciferase activity
0.75
0.50
0.25
0
****
***
NS
3xHA-LFY + EV
3xHA-LFY + UFO-3xFLAG
3xHA-LFY + UFOΔFbox-3xFLAG
3xHA-LFY + UFOdelF-3xFLAG
pAP3
3
2
1
3xHA-LFY + UFO-VP16-3xFLAG
3xHA-LFY + UFOΔFbox-VP16-3xFLAG
3xHA-LFY + UFOdelF-VP16-3xFLAG
3xHA-LFY-VP16 + EV
3xHA-LFY-VP16 + UFO-3xFLAG
3xHA-LFY-VP16 + UFOΔFbox-3xFLAG
3xHA-LFY-VP16 + UFOdelF-3xFLAG
D
35S::UFOΔFbox-5xmyc
35S::UFO-5xmyc
WT
Gain-of-function lines
Loss-of-function lines
#1 #2 #3 #4 #1 #2 #3
α-myc
35S::UFOΔFbox-3xFLAG
35S::UFO-3xFLAG
#1 #2 #3 #1 #2 #3 #4
α-FLAG
E
WT
ufo-1
35S::UFO-5xmyc ufo-1 (178-#19)
35S::UFOΔFbox-5xmyc ufo-1 (313-#2)
35S::UFO-3xFLAG ufo-1(177-#6)
35S::UFOΔFbox-3xFLAG ufo-1 (312-#2)
α-myc
α-FLAG
F
35S::UFO-5xmyc ufo-1 (line 178-#19)
35S::UFO-3xFLAG ufo-1 (line 177-#6)
35S::UFO-5xmyc ufo-1 (line 178-#2)
35S::UFO-3xFLAG ufo-1 (line 177-#2)
G
F-box domain
UFO (Arabidopsis thaliana) -------------MDSTVFINNPSLTLPFSYTFTSSSNSSTTTSTTTDSSSGQWMDGRIWSKLPPLLDRVIAFLPPPAFFRTRCVCKRFYSLLFSNTFL
DOT (Petunia hybrida) ----------MEAFHHAPISFHFPYAFPIPTPT-NFLGTTTPNSSSING---SPYMDSRLWSRLPHRLVDRIIAFLPPPAFFKARAVCKRFYGLLYSTNFL
AN (Solanum lycopersicum) ----------MEAFHHPPISFHFPYAFPIPTPTTNFLGT--PNSSSVNGMIINTWMDSRIWSRLPHRLIDRIIAFLPPPAFFRARVVCKRFYGLIYSTHFL
CsUFO (Cucumis sativus) ---MNFSHHMDMSLHSSMNPTIPFSYNIISSSNCGIISTTSNNNNVALLTTGPRMDSRIWSKLPQRILDRIVAFLPPPAFFRARCVCKRWYGLLFYASFL
STP (Pisum sativum) MEGFHPSMNMSMSMSMNMNPSFSYTFPITATASGAITNITTTTTTYNTTSTTPWMNSRIWSKLPHRLLDRIIAFLPPPAFFRARAVCKRFYSLLFSNSFL
PFO (Lotus japonicus) MEGFHPSMTMASPFPYTF-PISAGGIGVGVVGVGSSNLTTPTPTATGTYNNTPWMNSRIWSKLPQKLLDRVIAFLPTPAFFRARSVCKRWYSLLFSNTFL
TfUFO (Torenia fournieri) ------------MDHPFHHAPISVPPLLPYAFASNFHSFGANSSSSAGYTATDPFMDCRIWSRLPQRLVDRIIACLPPPAMFRARAVCKRWYALIFSTAFL
FIM (Antirrhinum majus) -------------MEAFQTPIFNLPL-PYGFTTTPNTI--NLQNTMIMSTTNPWMDCRIWSRLPQKLIDRIIACLPPPAFFRSRSVCKRWYSLIFSTTFL
GhUFO (Gerbera hybrida) ----------MEAFDHTHYVTTFPFSYPF-----------------STTTTTGNPSWMDSRIWSRLPQRLIDRVIAFLPPPAFFRARSVCKRWYSLLFSHSFL
CbUFO (Cosmos bipinnatus) ----------MEAFDHTHFVPTFPFPYPF-----------------ITTTTT-NPPWMDPRVWSRLPQRLLDRVIAFLPPPAFFRARSVCKRWYSLLFSHTFL
AqUFO1 (Aquilegia coerulea) ---------------------------MEQHINMFTNIPSSSNTYTARIGSPRMDPRIWSRLPDRLLARVICFLPPPAFFRFRSISDGWNRLLSSKCFV
APO1 (Oryza sativa) ------------------------------MMNPRRLPPLPSSTSSASAADDMDPRVWRRLPQPLVDRILACLPTPSFLRLRAACRRFYHLLFSSPFL

**Figure S1. UFO has SCF-dependent and independent functions, related to Figure 1**

(A to C) *pAP3* activation measured by DLRA in Arabidopsis protoplasts. EV = Empty Vector (pRT104-3xHA). UFOΔFbox corresponds to a deletion of the whole N-terminal part comprising the F-box domain (aa. 1-90), while UFOdelF corresponds to an internal deletion in the F-box domain (aa. 50-62; Risseeuw et al., 2013). Data represent averages of independent biological replicates and are presented as mean ± SD, each dot representing one biological replicate (n = 4). One-way ANOVA with Tukey's multiple comparisons test. Stars above bars represent a significant statistical difference compared to 3xHA-LFY + EV or 3xHA-LFY-VP16 + EV negative controls (NS: $p > 0.05$,*: $p < 0.05$, **: $p < 0.01$, ***: $p < 0.001$ and ****: $p < 0.0001$).

(D) Western Blot on protein extracts from T1 plants from different phenotypic classes described in Figure 1F. *35S::UFO-5xmyc* (178-#19) and *35S::UFO-3xFLAG* (177-#6) plants were used as positive controls. Total proteins were extracted from rosette leaves. Note the difference of molecular weight between UFO and UFOΔFbox. Loss-of-function defects are likely due to silencing of both transgene-encoded *UFOΔFbox* and endogenous *UFO.* Gels were cropped (see Supplemental Item 1).

(E) Western Blot on protein extracts from F2 plants described in Figure 1H and S1F. Total proteins were extracted from rosette leaves. Gels were cropped (see Supplemental Item 1).

(F) *ufo-1* complementation assay with *35S::UFO* lines. Rosette leaves (right, scale bar, 1 cm), inflorescence (middle, scale bar 1 mm) and flower (right, scale bar, 0.5 mm) phenotypes are shown. Primary inflorescences were removed to observe rosette phenotype. For each construct, two lines were crossed to *ufo-1* and at least 5 plants were analyzed per line. As in Risseeuw et al, our *35S::UFO* lines displayed relatively milder phenotypes than the *35S::UFO* phenotypes reported by Lee et al. (Lee et al., 1997; Risseeuw et al., 2013). Note that the *35S::UFO-5xmyc* 178-#2 line did not display the serrated leaves phenotype.

(G) Sequence alignment of UFO N-terminal region. The F-box domain is represented (Gagne et al., 2002). In selected species, presented proteins were identified as *UFO* homologs and their role was confirmed genetically (Chen et al., 2021; Ikeda et al., 2005; Levin and Meyerowitz, 1995; Li et al., 2019; Lippman et al., 2008; Sasaki et al., 2012; Sharma et al., 2019; Simon et al., 1994; Souer et al., 2008; Taylor et al., 2001; Zhang et al., 2003; Zhao et al., 2016).

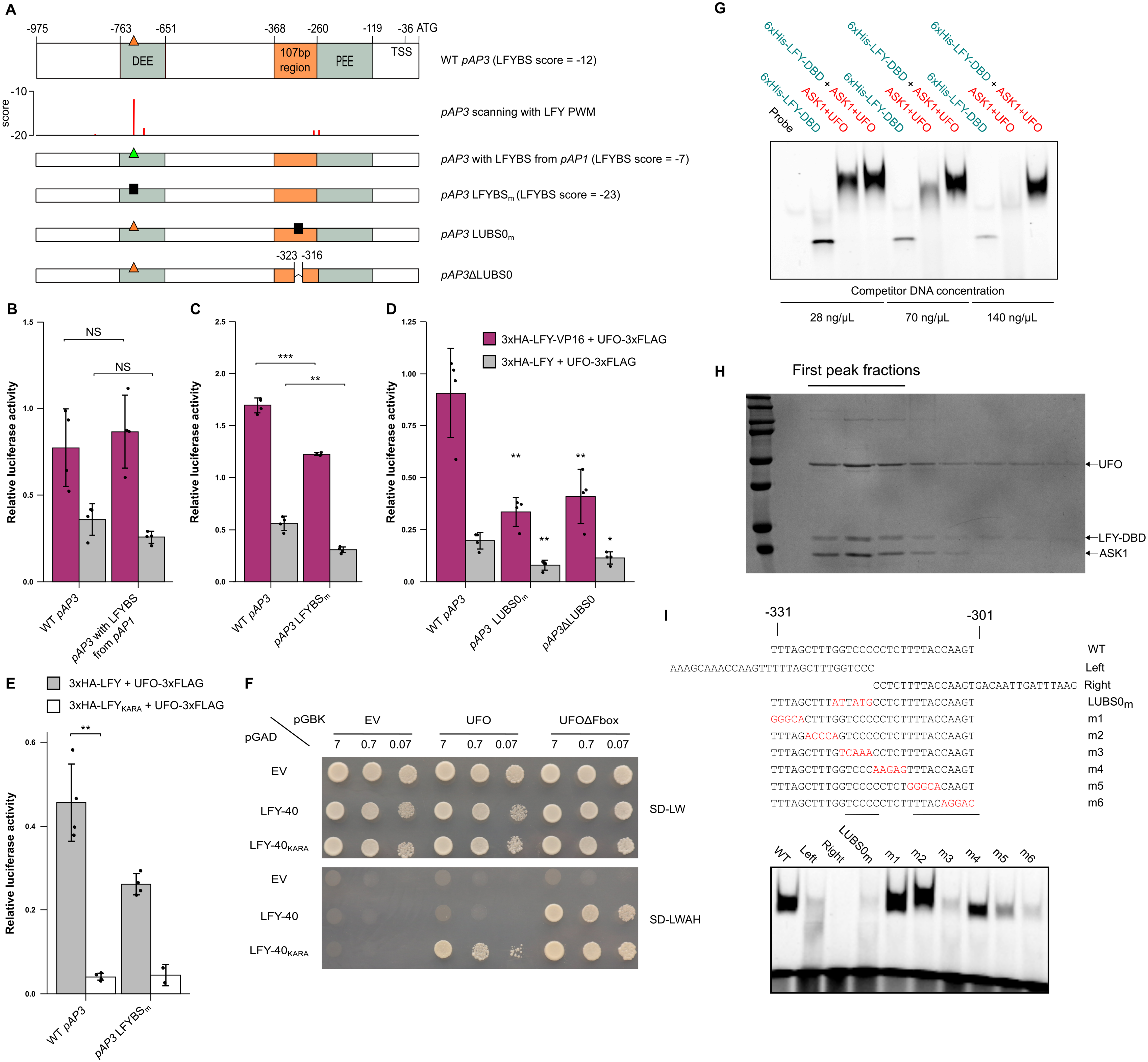

A
-975
-763
-651
-368
-260
-119
-36 ATG
DEE
107bp region
PEE
TSS
WT pAP3 (LFYBS score = -12)
score
-10
-20
pAP3 scanning with LFY PWM
pAP3 with LFYBS from pAP1 (LFYBS score = -7)
pAP3 LFYBSm (LFYBS score = -23)
pAP3 LUBS0m
-323 -316
pAP3ΔLUBS0
B
Relative luciferase activity
NS
NS
WT pAP3
pAP3 with LFYBS from pAP1
C
***
**
WT pAP3
pAP3 LFYBSm
D
3xHA-LFY-VP16 + UFO-3xFLAG
3xHA-LFY + UFO-3xFLAG
**
**
**
*
WT pAP3
pAP3 LUBS0m
pAP3ΔLUBS0
E
3xHA-LFY + UFO-3xFLAG
3xHA-LFYKARA + UFO-3xFLAG
**
WT pAP3
pAP3 LFYBSm
F
pGBK
pGAD
EV
UFO
UFOΔFbox
7
0.7
0.07
EV
LFY-40
LFY-40KARA
SD-LW
SD-LWAH
G
Probe
6xHis-LFY-DBD
ASK1+UFO
6xHis-LFY-DBD + ASK1+UFO
Competitor DNA concentration
28 ng/μL
70 ng/μL
140 ng/μL
H
First peak fractions
UFO
LFY-DBD
ASK1
I
-331
-301
TTTAGCTTTGGTCCCCCTCTTTTACCAAGT WT
AAAGCAAACCAAGTTTTTAGCTTTGGTCCC Left
CCTCTTTTACCAAGTGACAATTGATTTAAG Right
TTTAGCTTTATTATGCCTCTTTTACCAAGT LUBS0m
GGGCACTTTGGTCCCCCTCTTTTACCAAGT m1
TTTAGACCCAGTCCCCCTCTTTTACCAAGT m2
TTTAGCTTTGTCAAACCTCTTTTACCAAGT m3
TTTAGCTTTGGTCCCAAGAGTTTACCAAGT m4
TTTAGCTTTGGTCCCCCTCTGGGCACAAGT m5
TTTAGCTTTGGTCCCCCTCTTTTACAGGAC m6
WT
Left
Right
LUBS0m
m1
m2
m3
m4
m5
m6

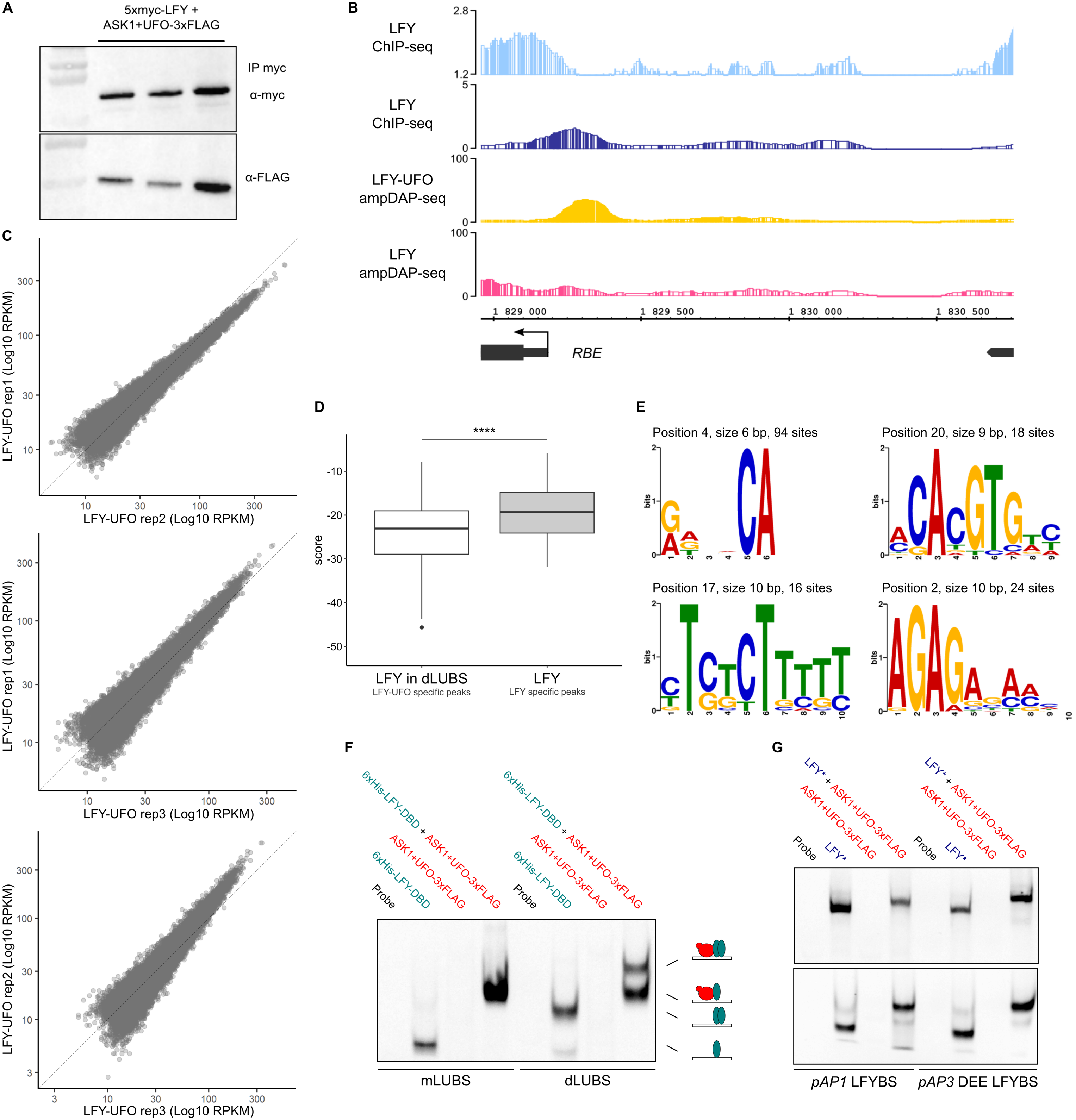
A
5xmyc-LFY +
ASK1+UFO-3xFLAG
IP myc
α-myc
α-FLAG
B
LFY
ChIP-seq
LFY
ChIP-seq
LFY-UFO
ampDAP-seq
LFY
ampDAP-seq
2.8
1.2
5
0
100
0
100
0
1 829 000
1 829 500
1 830 000
1 830 500
RBE
C
LFY-UFO rep1 (Log10 RPKM)
LFY-UFO rep2 (Log10 RPKM)
LFY-UFO rep1 (Log10 RPKM)
LFY-UFO rep3 (Log10 RPKM)
LFY-UFO rep2 (Log10 RPKM)
LFY-UFO rep3 (Log10 RPKM)
3
10
30
100
300
D
****
score
-10
-20
-30
-40
-50
LFY in dLUBS
LFY-UFO specific peaks
LFY
LFY specific peaks
E
Position 4, size 6 bp, 94 sites
Position 20, size 9 bp, 18 sites
Position 17, size 10 bp, 16 sites
Position 2, size 10 bp, 24 sites
bits
F
Probe
6xHis-LFY-DBD
ASK1+UFO-3xFLAG
6xHis-LFY-DBD + ASK1+UFO-3xFLAG
mLUBS
dLUBS
G
Probe
LFY*
ASK1+UFO-3xFLAG
LFY* + ASK1+UFO-3xFLAG
pAP1 LFYBS
pAP3 DEE LFYBS

**Figure S3. Genome-wide analysis of LFY-UFO DNA binding, related to Figure 3**

(A) Western Blot after DNA elution during ampDAP-seq experiment. After DNA elution, 20 µL of 1X SDS-PAGE Protein Sample Buffer was added to the remaining beads to run WB. Each lane represents one replicate. Gels were cropped (see Supplemental Item 1).

(B) IGB view of *pRBE* showing LFY ChIP-seq in inflorescences (light blue; Goslin et al., 2017) or seedlings (dark blue; Sayou et al., 2016), LFY-UFO ampDAP-seq (yellow; this study), LFY ampDAP-seq (pink; Lai et al., 2021), numbers indicate read number range.

(C) Assessment of experimental reproducibility of ampDAP-seq experiment through the comparison of replicates datasets 2 by 2.

(D) Score distribution of LFY PWM with dependencies (Moyroud et al., 2011) within dLUBS (best site on 20% most LFY-UFO-specific genomic regions, high CFC) and in canonical LFYBS (best site on 20% most LFY-specific genomic regions, low CFC). Best sites were selected within ±25 bp around the peak maximum (see Methods). Wilcoxon rank sum test (****: $p < 0.0001$).

(E) *De novo* identification of URM from LFY ChIP-seq data. The panel shows the motifs identified at a fixed distance from LFY canonical binding sites in 298 regions harboring high LFY ChIP-seq to LFY ampDAP-seq coverage ratio. The text above each motif gives the motif's start position relative to the canonical LFYBS, its length and the number of sites used to build the motif.

(F) EMSA with mLUBS and dLUBS highest score sequences. 6xHis-LFY-DBD and ASK1-UFO-3xFLAG complex are recombinant. Drawings represent the different types of complexes involving LFY-DBD (pale blue) and ASK1-UFO (red) on DNA. LFY-DBD binds as a monomer as previously reported (Hamès et al., 2008). Gel was cropped and only protein-DNA complexes are shown (see Supplemental Item 1).

(G) EMSA with DNA probes corresponding to *pAP1* and *pAP3* DEE LFYBS. LFY* refers either to *in vitro*-produced 5xmyc-LFY (top) or recombinant 6xHis-LFY-DBD (bottom). Note that probes used here have the same length as those used to study LUBS. Gels were cropped and only protein-DNA complexes are shown. Gels were cropped and only protein-DNA complexes are shown (see Supplemental item 1).

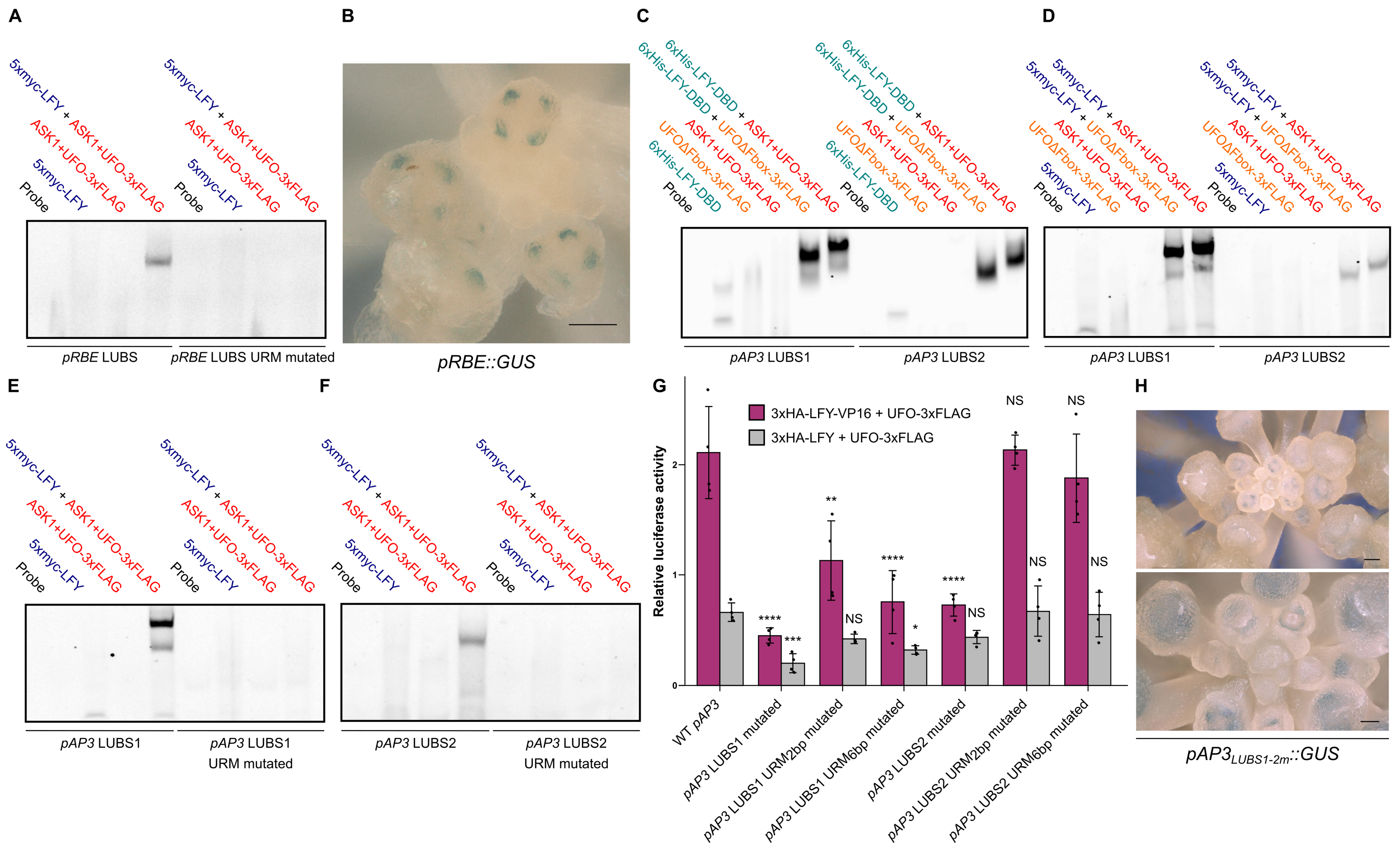


I

| Gene | Name | Description | Up/Down regulated | LFY ChIP-Seq set |
|---|---|---|---|---|
| AT1G13260 | RAV1 | related to ABI3/VP1 1 | down | A |
| AT1G24530 | NA | Transducin/WD40 repeat-like superfamily protein | down | B |
| AT1G34110 | NA | Leucine-rich receptor-like protein kinase family protein | up | C; B |
| AT1G54010 | NA | GDSL-like Lipase/Acylhydrolase superfamily protein | down | A |
| AT1G73805 | NA | Calmodulin binding protein-like | down | C; B; A |
| AT1G74440 | NA | Protein of unknown function (DUF962) | down | A |
| AT1G75450 | ATCKX5 | cytokinin oxidase 5 | up | C; B; D |
| AT1G76110 | NA | HMG (high mobility group) box protein with ARID/BRIGHT DNA-binding domain | up | A |
| AT1G80840 | ATWRKY40 | WRKY DNA-binding protein 40 | up | C; B; A |
| AT3G15270 | SPL5 | squamosa promoter binding protein-like 5 | up | A |
| AT3G20810 | NA | 2-oxoglutarate (2OG) and Fe(II)-dependent oxygenase superfamily protein | up | A |
| AT3G28180 | ATCSLC04 | Cellulose-synthase-like C4 | down | B |
| AT3G54340 | AP3 | K-box region and MADS-box transcription factor family protein | up | B |
| AT3G55560 | AGF2 | AT-hook protein of GA feedback 2 | up | C; B; A |
| AT4G00730 | ANL2 | Homeobox-leucine zipper family protein / lipid-binding START domain-containing protein | up | A |
| AT4G02380 | AtLEA5 | senescence-associated gene 21 | down | B |
| AT4G34160 | CYCD3 | CYCLIN D3 | up | A |
| AT4G35900 | FD | Basic-leucine zipper (bZIP) transcription factor family protein | down | C; B; A |
| AT5G02540 | NA | NAD(P)-binding Rossmann-fold superfamily protein | up | B; D |
| AT5G20240 | PI | K-box region and MADS-box transcription factor family protein | up | A |

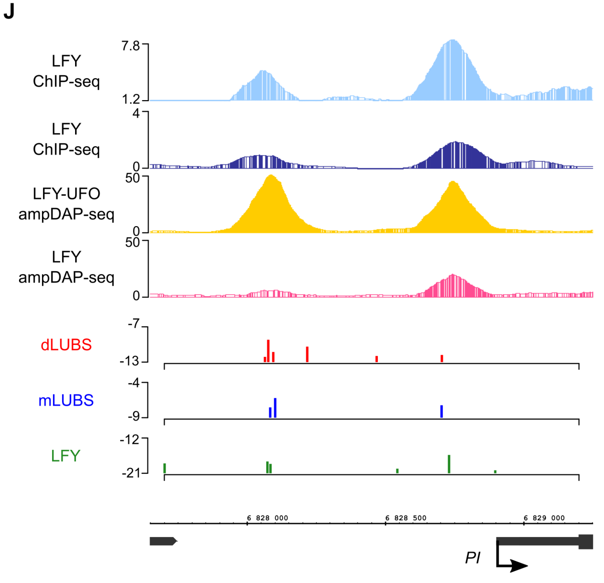

**Figure S4. Characterization of *pRBE* and *pAP3* LUBS, related to Figure 4**

(A) EMSA with probes corresponding to *pRBE* LUBS*,* WT or with URM mutated.

(B) *In vivo* analysis of *pRBE::GUS* fusions. Same as in Figure 4C, with another view showing staining in the four petal primordia (scale bar, 50 µm).

(C-D) EMSA with *pAP3* LUBS1 and LUBS2 DNA probes and recombinant 6xHis-LFY-DBD (C) or *in vitro*-produced 5xmyc-LFY (D). Note the difference of complex size between UFO and UFOΔFbox.

(E-F) EMSA with DNA probes corresponding to *pAP3* LUBS1 (E) and LUBS2 (F), WT or with URM mutated.

(G) Promoter activation measured by DLRA in Arabidopsis protoplasts with indicated effectors. Different promoter versions were tested as indicated under x-axis. Either 2 bp (high-informative CA) or 6 bp (whole URM) of *pAP3* LUBS1 and LUBS2 URM were mutated. Data represent averages of independent biological replicates and are presented as mean ± SD, each dot representing one biological replicate (n = 4). One-way ANOVA with Tukey's multiple comparisons tests. One-way ANOVA were performed with data from the same effector and stars represent a statistical difference compared to WT *pAP3* promoter. (NS: p > 0.05,*: p < 0.05, **: p < 0.01, ***: p < 0.001 and ****: p < 0.0001).

(H) *In vivo* analysis of $pAP3_{LUBS1\text{-}2m}$*::GUS* fusions. Same as in Figure 4F, except that staining incubation time was increased to 17 h. Representative pictures are shown (top scale bar, 100 µm, bottom scale bar, 50 µm). The faint *AP3* pattern suggests that other LUBS (such as LUBS0) may take over but less efficiently. Note that with this staining incubation time, all plants expressing *pAP3::GUS* showed a highly saturated staining.

(I) List of candidate LFY-UFO target genes selected as i) present in regions specifically bound by LFY-UFO in ampDAP-seq (high CFC) ii) bound *in vivo* in LFY ChIP-seq experiments (Goslin et al., 2017 (A); Sayou et al., 2016 (B); Moyroud et al., 2011 (C); Jin et al., 2021 (D)) and iii) deregulated in *ufo* inflorescences (Schmid et al., 2005).

(J) From the top: IGB view of *PISTILLATA* promoter region showing LFY ChIP-seq in inflorescences (light blue; Goslin et al., 2017) or seedlings (dark blue; Sayou et al., 2016), LFY-UFO ampDAP-seq (yellow; this study), LFY ampDAP-seq (pink; Lai et al., 2021), numbers indicate read number range. Below, predicted binding sites using the dLUBS, mLUBS models from Figure 3C and LFY PWM with dependencies (Moyroud et al., 2011), y-axis represents score values.

For all EMSAs, gel pictures were cropped and only protein-DNA complexes are shown (see Supplemental Item 1).

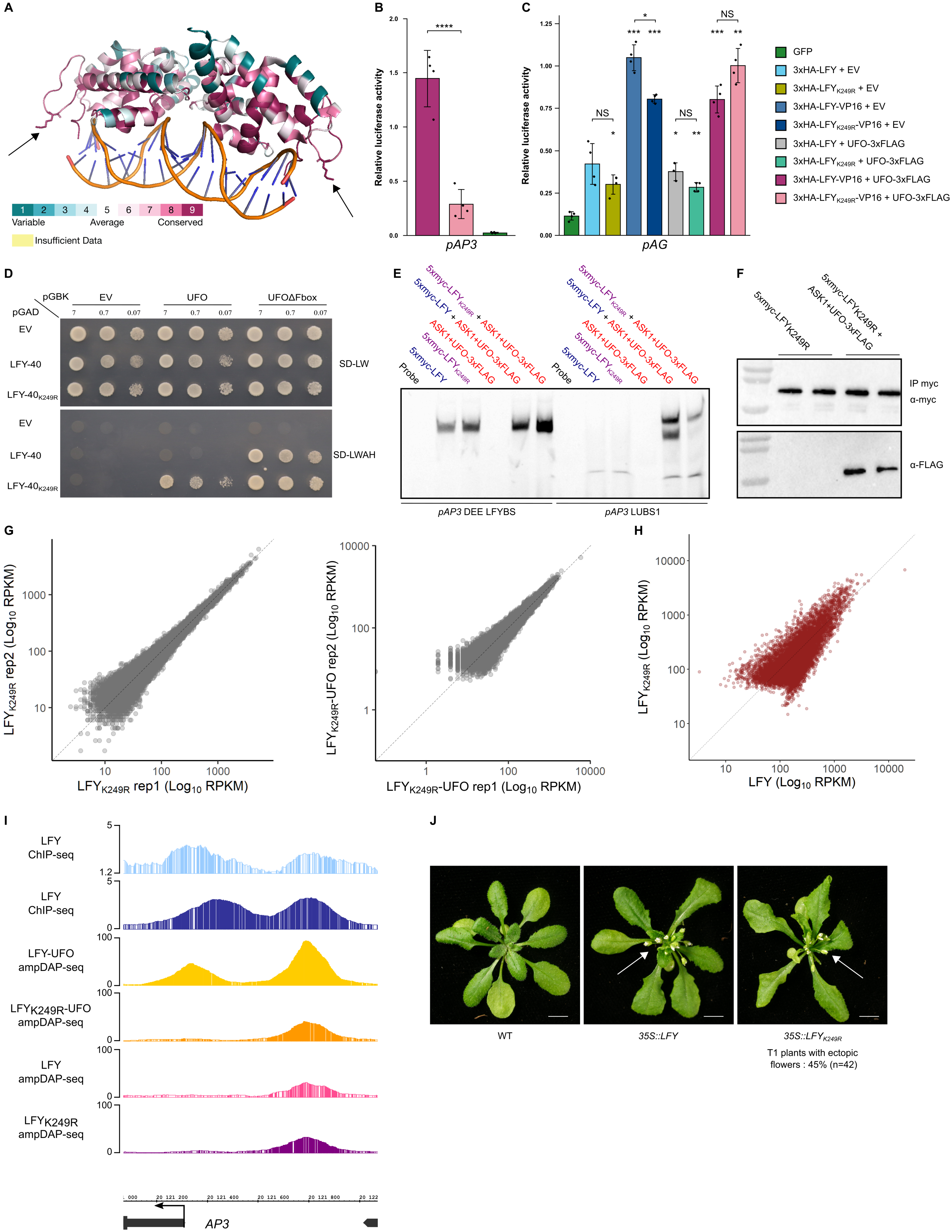

A
Variable
Average
Conserved
Insufficient Data
B
Relative luciferase activity
pAP3
C
pAG
GFP
3xHA-LFY + EV
3xHA-LFYK249R + EV
3xHA-LFY-VP16 + EV
3xHA-LFYK249R-VP16 + EV
3xHA-LFY + UFO-3xFLAG
3xHA-LFYK249R + UFO-3xFLAG
3xHA-LFY-VP16 + UFO-3xFLAG
3xHA-LFYK249R-VP16 + UFO-3xFLAG
D
pGBK
pGAD
EV
UFO
UFOΔFbox
LFY-40
LFY-40K249R
SD-LW
SD-LWAH
E
Probe
5xmyc-LFY
5xmyc-LFYK249R
ASK1+UFO-3xFLAG
5xmyc-LFY + ASK1+UFO-3xFLAG
5xmyc-LFYK249R + ASK1+UFO-3xFLAG
pAP3 DEE LFYBS
pAP3 LUBS1
F
5xmyc-LFYK249R
5xmyc-LFYK249R + ASK1+UFO-3xFLAG
IP myc
α-myc
α-FLAG
G
LFYK249R rep2 (Log10 RPKM)
LFYK249R rep1 (Log10 RPKM)
LFYK249R-UFO rep2 (Log10 RPKM)
LFYK249R-UFO rep1 (Log10 RPKM)
H
LFYK249R (Log10 RPKM)
LFY (Log10 RPKM)
I
LFY ChIP-seq
LFY ChIP-seq
LFY-UFO ampDAP-seq
LFYK249R-UFO ampDAP-seq
LFY ampDAP-seq
LFYK249R ampDAP-seq
AP3
J
WT
35S::LFY
35S::LFYK249R
T1 plants with ectopic flowers : 45% (n=42)

**Figure S5. The LFY K249 is essential for LFY-UFO-LUBS complex formation, related to Figure 5**

(A) Structure of LFY-DBD (Hamès et al., 2008). Residues were colored by conservation using Consurf with default parameters (Ashkenazy et al., 2016). K249 residues on each LFY monomer are represented as sticks and indicated with arrows. Note that the K249-containing loop is highly conserved.

(B-C) Promoter activation measured by DLRA in Arabidopsis protoplasts with indicated effectors (right). EV = Empty Vector (pRT104-3xHA). Tested promoters are indicated below each graph. Note that for 3xHA-LFY+UFO-3xFLAG on *pAG* only n = 3 biological replicates are shown. Data represent averages of independent biological replicates and are presented as mean ± SD, each dot representing one biological replicate (n = 4 unless specified). One-way ANOVA with Tukey's multiple comparisons tests (B) or Welch's ANOVA with Games-Howell post-hoc test (C). In (C), stars above bars represent a statistical difference compared to GFP. (NS: $p > 0.05$,*: $p < 0.05$, **: $p < 0.01$, ***: $p < 0.001$ and ****: $p < 0.0001$)

(D) Effect of the $LFY_{K249R}$ mutation on LFY-UFO interaction in Y2H. EV = Empty Vector. LFY-40 is a LFY version lacking the first 40 aa and better tolerated by yeast cells. Values correspond to the different dilutions (OD = 7, 0.7 and 0.07). Top picture corresponds to the non-selective plate lacking Leucine and Tryptophan (SD -L-W), and bottom picture corresponds to the selective plate lacking Leucine, Tryptophan, Histidine and Adenine (SD -L-W-A-H). Pictures were taken at day + 4.

(E) EMSA with DNA probes corresponding to *pAP3* DEE LFYBS and *pAP3* LUBS1 and indicated proteins. *pAP3* DEE LFYBS DNA probe was used as a control for binding on canonical LFYBS. Gel was cropped and only protein-DNA complexes are shown (see Supplemental Item 1).

(F) WB after DNA elution during ampDAP-seq experiment. After DNA elution, 20 µL of 1X SDS-PAGE Protein Sample Buffer was added to the remaining beads to run WB. Each lane represents one replicate. Gels were cropped (see Supplemental Item 1).

(G) Reproducibility of ampDAP-seq experiments with $LFY_{K249R}$ (left) and $LFY_{K249R}$-UFO (right) through the comparison of replicates datasets 2 by 2.

(H) Comparison of peak coverage in $LFY_{K249R}$ (y-axis, this study) and LFY (x-axis, Lai et al., 2021) ampDAP-seq experiments.

(I) Integrated Genome Browser (IGB) view of *pAP3* showing LFY ChIP-seq in inflorescences (light blue; Goslin et al., 2017) or seedlings (dark blue; Sayou et al., 2016), LFY-UFO ampDAP-seq (yellow; this study), $LFY_{K249R}$-UFO ampDAP-seq (orange; this study), LFY ampDAP-seq (pink; Lai et al., 2021), and $LFY_{K249R}$ ampDAP-seq (purple; this study). Numbers indicate read number range.

(J) Pictures of WT and representative transgenic plants expressing *35S::LFY* or *35S::*$LFY_{K249R}$ (scale bar, 1 cm). The white arrows indicate ectopic rosette flowers. *35S::LFY* was obtained previously (Sayou et al., 2016). 42 T1 plants expressing *35S::*$LFY_{K249R}$ were analyzed; the percentage of plants with a LFY overexpressing phenotype is comparable to the one obtained with *35S::LFY* (Sayou et al., 2016).

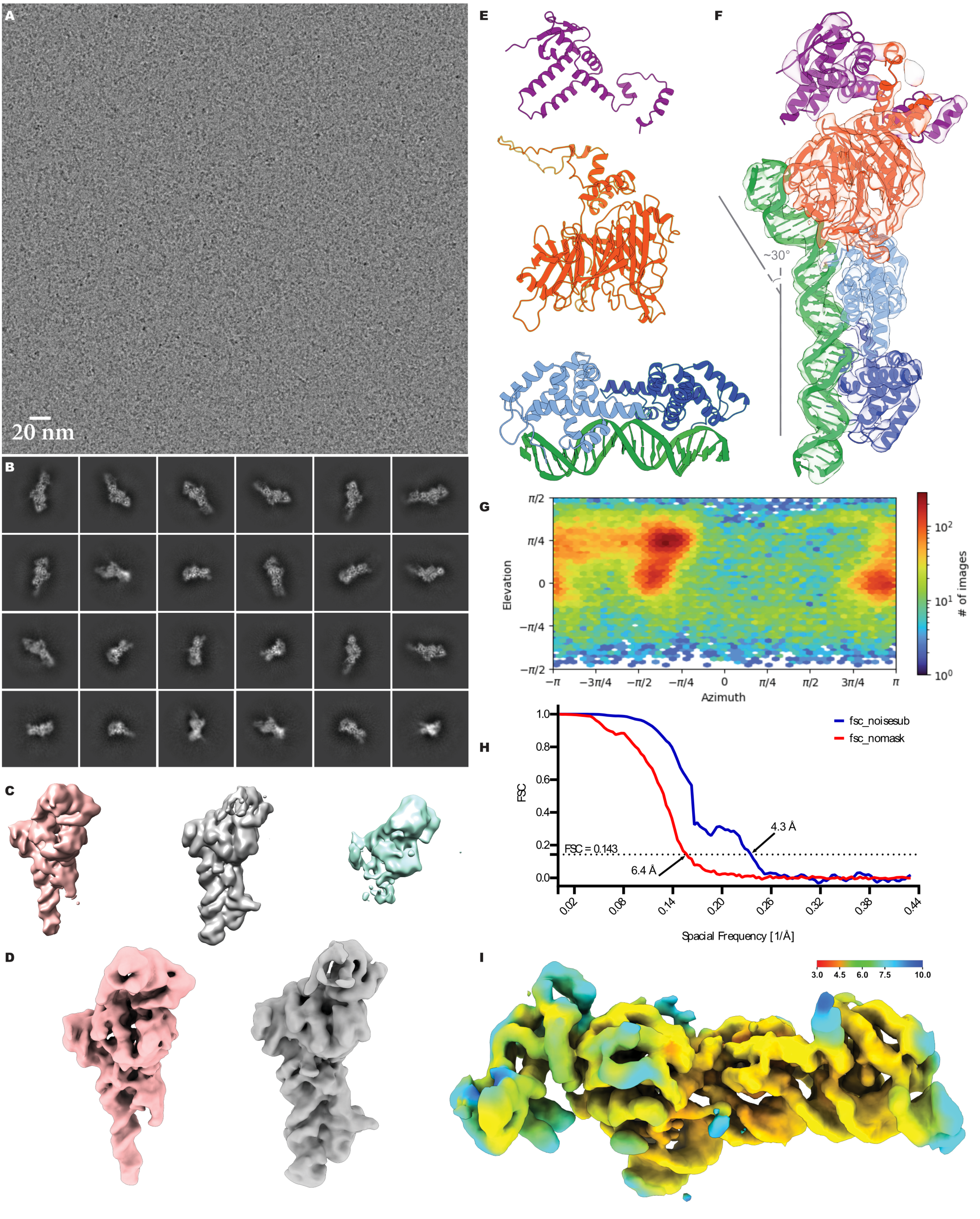
A
20 nm
B
C
D
E
F
~30°
G
π/2
π/4
0
−π/4
−π/2
Elevation
−π
−3π/4
−π/2
−π/4
0
π/4
π/2
3π/4
π
Azimuth
$10^2$
$10^1$
$10^0$
### of images
H
fsc_noisesub
fsc_nomask
FSC
1.0
0.8
0.6
0.4
0.2
0.0
4.3 Å
FSC = 0.143
6.4 Å
0.02
0.08
0.14
0.20
0.26
0.32
0.38
0.44
Spacial Frequency [1/Å]
I
3.0
4.5
6.0
7.5
10.0

**Figure S6. UFO binds DNA and LFY, related to Figure 6**

(A) A representative micrograph of the ASK1-UFO-LFY-DNA complex in vitreous ice.

(B) Selected 2D class averages of the particles submitted to *ab initio* reconstruction and heterogeneous refinement for 3D classification.

(C) Intermediate reconstructions of the 3D classes after heterogeneous refinement.

(D) Final reconstructions of ASK1-UFO-LFY-DNA complexes (involving either a LFY-DBD monomer (pink) or a LFY-DBD dimer (gray)) after Non-Uniform refinement.

(E) Unprocessed AlphaFold2 model for ASK1 (top, purple; uniprot ID, Q39255), UFO (middle, red; uniprot ID, Q39090) and the LFY-DBD dimer/DNA crystallographic structure (bottom, pale and dark blue for the LFY-DBD dimer and green for the DNA; PDB entry: 2VY1).

(F) Cryo-EM density map color-coded by fitted molecule. Note the kink on DNA induced by the presence of UFO.

(G) Heat map of the angular distribution of particle projections contributing for the final reconstruction of the complete ASK1-UFO-LFY-DNA complex (with a LFY-DBD dimer).

(H) Gold-standard Fourier shell correlation (FSC) curves. The dotted line represents the 0.143 FSC threshold, which indicates a nominal resolution of 6.4 Å for the unmasked (red) and 4.3 Å for the masked (blue) reconstruction.

(I) View of the post-processed map of the complete ASK1-UFO-LFY-DNA complex, colored according to the local resolution.
